# Sleep arousals are associated with the polygenic risk for developing Alzheimer’s disease and with cognitive decline in healthy late middle-aged individuals

**DOI:** 10.1101/2025.11.11.687823

**Authors:** Nasrin Mortazavi, Mikhail Zubkov, Daphne Chylinski, Fabienne Collette, Christine Bastin, Pierre Maquet, Gilles Vandewalle, Puneet Talwar

**Affiliations:** GIGA-Institute, CRC-Human Imaging, University of Liège, Belgium; Psychology and Cognitive Neuroscience Research Unit, University of Liège, Belgium; Department of Neurology, CHU of Liège, Liège, Belgium

**Keywords:** Alzheimer’s disease, arousal, Polygenic risk score, sleep

## Abstract

**Objective:** Sleep disturbances are increasingly recognized as early features of Alzheimer’s disease (AD) neuropathology. In that context, spontaneous arousals during sleep have been associated with the burden of Amyloid beta in the brain of healthy late middle-aged individuals. Whether the heterogeneity of arousals during sleep may be related to the genetic risk of developing AD in young adults is not established. Likewise, whether arousals may be associated with cognitive decline is not known. Here, we evaluated the association between arousals, the genetic risk for developing AD and cognitive performance and cognitive decline in healthy young and late-middle-aged individuals.

**Methods:** We classified spontaneous arousals using in-lab EEG recordings of sleep in 453 younger individuals (22+/-2.7y; 49 women) and 87 late middle-aged individuals (59.3+/-5.3y; 59 women) based on their association with sleep stage transitions and changes in muscle tone. We examined the associations between arousal types and the polygenic risk scores (PRS) for AD, cognitive performance at baseline and, in late middle-aged individuals, cognitive decline over 2 and 7 years.

**Results:** The prevalence of arousals associated with sleep stage transition was higher in late middle-aged vs. younger individuals. Among these arousals, those with and without muscle tone increases were, respectively, associated with lower and higher PRS for AD in late middle-aged but not in younger individuals. In the late middle-aged individuals, transition arousals associated with and without muscle tone increases were, respectively, correlated with better and worse attentional performance at baseline, and lower and larger memory decline over 2 or 7 years.

**Conclusion:** The heterogeneity in spontaneous arousals during sleep may reflect their physiological intensity or underlying neural activation, and may indicate vulnerability to AD in late middle-aged individuals. The findings may contribute to identifying early markers of neurodegenerative risk.

**Statement of Significance:** Sleep arousals are typically regarded as disruptive events, yet their physiological diversity may reveal important insights into brain health. In this study, we report that distinct subtypes of spontaneous sleep arousals are differentially associated with genetic vulnerability to Alzheimer’s disease (AD) and with future cognitive decline in healthy late middle-aged adults. Specifically, sleep arousals linked to sleep stage transitions but lacking muscle activation were related to higher polygenic risk for AD and greater memory decline, while those accompanied by muscle tone increases showed the opposite pattern. These findings indicate that subtle variations in sleep microstructure can reflect neurobiological vulnerability to AD before clinical symptoms emerge. By identifying electrophysiological markers associated with genetic risk and cognitive trajectories, this work advances the potential for using sleep-based biomarkers to detect and monitor preclinical neurodegenerative processes.

## Introduction

Individuals with Alzheimer’s disease (AD) — who are characterized by cognitive deficits, particularly in memory — also show decreased total sleep time, sleep efficiency, N3 sleep and REM sleep, and increased number of awakenings compared with controls (Zhang et al., 2022) and these changes have been correlated with the severity of their cognitive impairment (Zhang et al., 2022). Alterations in sleep are also related to the hallmarks of AD pathogenesis (i.e., Amyloid-Beta (Aβ) and tau accumulation) in healthy individuals, prior to any clinical manifestations of the disease (Van Egroo et al., 2019). In addition, sleep disturbances have been linked to the future risk of both cognitive decline and AD pathology (Shi et al., 2018). Notably, individuals at higher risk for AD exhibit more frequent nocturnal arousals – transient/abrupt shift of EEG frequency not associated with full awakenings– than those at lower risk (Tsai et al., 2022). Additionally, arousals related to movement are associated with increased risk of developing AD and cognitive decline (Lim et al., 2013). In cognitively healthy late middle-aged individuals, sleep arousals were associated with early Aβ burden depending on whether arousals were linked to sleep stage transitions (T+/T–) and to detectable increase in muscle tone (M+/M−) (Chylinski et al., 2021). Specifically, T+M– arousals were associated with higher Aβ burden while T–M+ arousals were linked to lower Aβ accumulation and better cognitive performance, particularly in the attentional domain.

Recent studies in rodents indicate that the heterogeneity in arousals may be related to the activity of locus coeruleus (LC), which is among the first sites of tau protein aggregation in the brain (Braak & Del Tredici, 2012) and is likely to contribute to the alteration in sleep regulation found in preclinical AD (Van Egroo et al., 2024). Stronger activity of the LC during slow wave sleep would be associated with increased arousal density while lower level of LC activation would lead to decreased arousal density (Osorio-Forero et al., 2022). The heterogeneity in sleep arousal appears therefore to be directly related to a key structure for sleep and for AD, and as a result, may constitute a marker of the vulnerability for AD. How early this vulnerability can be detected through arousals and whether it bears prediction for future cognitive decline is unknown.

Aβ protein accumulation in the brain typically begins to increase over the sixth decade in humans (Fleisher et al., 2013) and is therefore not a useful biomarker for assessing AD risk in younger individuals. In contrast, genetic approaches offer unique tools to assess the variability for the risk for complex diseases, including AD. AD is recognized to be polygenic (Escott-Price et al., 2015; Talwar et al., 2016) and genome-wide association studies (GWAS) have identified over 70 genes which are associated with AD (Reitz et al., 2023). Due to the polygenic nature of AD, polygenic risk scores (PRS)—which aggregate the risks associated with common genetic variants— have proven to be an effective method for assessing AD risk. Studies have demonstrated that PRS can distinguish between AD cases and healthy controls, reaching a prediction accuracy up to between 75% and 84% (Escott-Price et al., 2015; Escott-Price et al., 2017). The PRS would be particularly useful for studying asymptomatic individuals of any age for studying AD risk long before classical pathological hallmarks are detectable (Baker & Escott-Price, 2020). Importantly, PRS may not grasp the same aspect of AD risk than Aβ assessment as reported in studies of cognitively healthy individuals where PRS was not associated with baseline Aβ burden but could predict future Aβ accumulation in longitudinal follow-up (Ge et al., 2018; Luckett et al., 2022; Xicota et al., 2022).

In this study, we examined the associations between PRS for AD, different types of sleep arousals—classified by the presence of muscle tone increase and sleep stage transitions—and cognitive performance in a relatively large sample of cognitively unimpaired individuals (N=540). Our aim was to determine whether arousal types previously associated with Aβ accumulation were also linked to PRS for AD in late middle-aged individuals aged 50 to 70y and in much younger adults aged 18 to 31y. We anticipated that more T+M− and T−M+ arousals would be linked, respectively, to higher and lower PRS values. We further assess their link to cognitive performance and, in late-middle aged individuals, with cognitive decline over 2y and 7y follow-up periods. We hypothesize that, in late middle-aged individuals, T−M+ arousals would be associated with better attentional performance in the baseline (as observed in our previous study (Chylinski et al., 2021)) while they would be associated with reduced memory decline at follow-up, given that memory deterioration is a core feature of AD.

## Methods

### Participants

A total of 540 healthy participants [453 younger individuals aged 18–31 years (22.04 ± 2.69y; 49 women;) and 87 late middle-aged individuals aged 50-69 years (59.28 ± 5.3y; 59 women)] part of different multi-modal studies were included in the present analyses **(Table 1)** (Chylinski et al., 2021; Muto et al., 2021). Data from young adults were collected across 6 different studies while data from late middle-aged individuals were collected as part of a single study. One of the studies in young adults contributed the majority of the sample, and including 357 Caucasian young men (to reduce variability as the study was originally designed as a genetic study). This means that vast majority of our young sample consist of men (~90%) while a minority of men constituted the late-middle aged sample (~32%). This constitutes an inherent limitation of the present study, even if sex is included as covariates in all statistical analyses. All studies collected quantitative sleep parameters and blood samples or buccal swabs to assess PRS for AD (**Figure 1**). Extensive cognitive evaluation was conducted in the largest studies of the younger dataset (N=357) and in all late middle-aged individuals, while the later part of the sample also completed follow-up cognitive assessment after 2 and 7 years.

**Figure 1:**
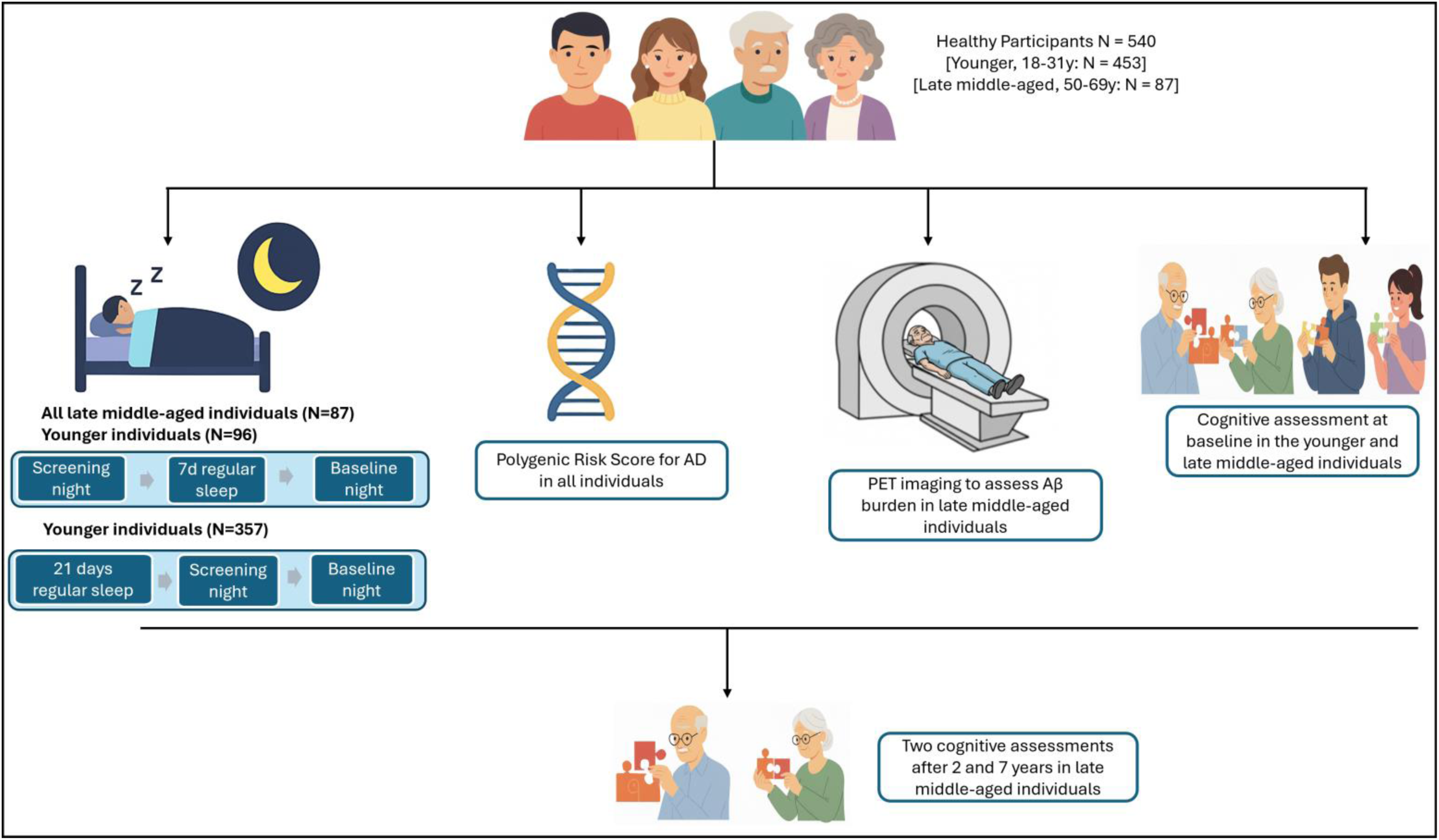
Overview of the study design. The habitual sleep of young and older adults was recorded in a laboratory setting to identify sleep arousals. Blood samples or buccal swabs were then collected to measure Alzheimer’s disease polygenic risk scores (PRS), and PET imaging was conducted in late middle-aged individuals to assess amyloid-beta (Aβ) burden. Participants subsequently completed a battery of cognitive tests. A subset of late middle-aged adults who initially participated in the study also returned for follow-up cognitive assessments after 2 and 7 years.

**Table 1.** Characteristics of the study sample. The p-values shown in the table correspond to two-sample t-tests except for sex that were compared using a Chi-square test. Sleep quality was assessed by the Pittsburgh Sleep Quality index (PSQI) (Buysse et al., 1989). Daytime sleepiness was measured by the Epworth Sleepiness Scale (Johns, 1991), Chronotype was assessed by the Morningness-Eveningness Questionnaire (MEQ) (Horne & Ostberg, 1976). Anxiety was estimated by the Beck Anxiety Inventory (Beck, Epstein, et al., 1988) and Depression was estimated by the 21-item Beck Depression Inventory II (Beck, Steer, et al., 1988); Total sleep time was extracted from polysomnography recordings. All late–middle-aged individuals and the largest young adult dataset (N=357) had measures of anxiety, depression, sleep quality, daytime sleepiness, and chronotype. These measures were not available for the remaining young adult datasets (N=96). Women were largely under-represented in the younger group, due to the selection criteria of one of the studies composing the sample (see methods), while they represent a large majority of the late-middle-aged group. This constitutes an inherent limitation of the present study, even if sex is included as covariates in all statistical analyses.

| Characteristic | All<br>Mean (SD) | Young<br>Mean (SD) | Late middle-aged - Mean<br>(SD) | p-value |
| --- | --- | --- | --- | --- |
| Sample size (N) | 540 | 453 | 87 | - |
| Sex (Men) | 80.29% | 89.37% | 32.18% | <.0001 |
| Age (years) | 29.27 (15.07) | 22.04 (2.69) | 59.28 (5.35) | <.0001 |
| Body Mass Index (BMI) (kg*m<sup>-2</sup>) | 22.67 (2.64) | 22.15 (2.31) | 24.83 (2.85) | <.0001 |
| Anxiety | 2.60 (2.92) | 2.56 (2.94) | 2.78 (2.86) | 0.5 |
| Depression | 3.46 (3.88) | 2.96 (3.53) | 5.52 (4.55) | <.0001 |
| Sleep Quality | 3.68 (1.99) | 3.46 (1.76) | 4.63 (2.59) | 0.0001 |
| Daytime sleepiness | 5.97 (3.63) | 5.94 (3.54) | 6.12 (4.00) | 0.67 |
| Chronotype | 50.80 (8.30) | 50.10 (8.26) | 53.68 (7.88) | 0.0003 |
| Total sleep time (min) | 439.9 (47.5) | 451.4 (41.53) | 392.3 (40.90) | <.0001 |
| T+M+ arousals number | 13.45 (5.88) | 12.92 (5.53) | 15.16 (6.57) | 0.004 |
| T+M- arousals number | 9.98 (5.79) | 9.50 (5.21) | 13.59 (7.29) | <.0001 |
| T-M+ arousals number | 96.12 (37.43) | 106.2 (33.12) | 55.60 (25.88) | <.0001 |
| T-M- arousals number | 118.64 (44.73) | 127.9 (45.59) | 95.71 (44.08) | <.0001 |
| T+M+ arousals density (number/h) | 1.82 (0.81) | 1.72 (0.72) | 2.33 (1.02) | <.0001 |
| T+M- arousals density (number/h) | 1.40 (0.87) | 1.27 (0.72) | 2.11 (1.19) | <.0001 |
| T-M+ arousals density (number/h) | 13.25 (4.62) | 14.16 (4.17) | 8.54 (3.98) | <.0001 |
| T-M- arousals density (number/h) | 16.70 (6.18) | 17.11 (6.07) | 14.59 (6.34) | 0.0005 |
| Aβ burden (centiloid units) | - | - | -5.19 (8.68) | - |

In all studies, participants were excluded if they had a body mass index (BMI) less than 18 and greater than 27 kg/m^2^ (greater than 29 kg/m^2^ for late middle-aged individuals), a history of psychiatric disorders or severe brain injury, documented/diagnosed sleep pathologies such as insomnia and REM behavior disorder, substance addiction, chronic use of medication affecting the central nervous system, smoking, excessive alcohol consumption (more than 14 units per week), high caffeine intake (more than three cups per day for younger participants, or more than five cups per day for older participants), shift work within the past 6 months, trans-meridian travel within the preceding two months, moderate to severe subjective anxiety or depression, as indicated by a score greater than 16 on the Beck Anxiety Inventory (BAI) (Beck, Epstein, et al., 1988) or greater than 19 on the Beck Depression Inventory-II (BDI-II) (Beck, Steer, et al., 1988). Poor sleep quality (Pittsburgh Sleep Quality Index score > 7) (Buysse et al., 1989), excessive daytime sleepiness (Epworth Sleepiness Scale score > 14) (Johns, 1991), significant sleep apnea (apnea-hypopnea index > 15 events per hour), and parasomnia as determined during an in-laboratory screening night using polysomnography (PSG) and based on the 2017 American Academy of Sleep Medicine (AASM) criteria (version 2.4) (Berry et al., 2012), also led to exclusion. Old individuals were also excluded if they had clinical symptoms of cognitive impairment (dementia rating scale < 130; mini mental state examination < 27).

All study procedures were approved by the Ethics Committee of the Faculty of Medicine at the University of Liège (Belgium). Written informed consent was obtained from all participants prior to inclusion and participants received financial compensation for their participation. The study was conducted in accordance with the Declaration of Helsinki and the World Medical Association’s International Code of Medical Ethics.

### Protocol

Data from young adults were obtained from six separate studies (Gaggioni et al., 2019; Koshmanova et al., 2022; Ly et al., 2016; Mascetti et al., 2013; Muto et al., 2016, 2021; Vandewalle et al., 2009), whereas data from late middle-aged individuals were derived from a single study (Chylinski et al., 2021) and we detail here only those aspects that are relevant for the present study.

For the majority of young participants (N=357 young men), for three weeks prior to the in-lab experiment, participants adhered to a regular sleep schedule based on their usual sleep times (within ±30 minutes for the first two weeks and within ±15 minutes for the final week), verified through actigraphy data (Actiwatch 4, CamNtech, Cambridge, UK). Prior to the experiment, participants completed a urine drug screen (Multipanel Drug Test, SureScreen Diagnostics Ltd) and an adaptation night in the laboratory aligned with their habitual sleep-wake schedule. During this night, full PSG recordings were performed to screen for sleep-related breathing disorders and periodic limb movements. On the second day, participants left the lab in the morning and were instructed not to nap during the day, with adherence confirmed via actigraphy. They returned to the lab in the evening, 3.5 hours before their scheduled lights-off time, and had a baseline night of sleep recorded in complete darkness. The timing of this night was centered on the average sleep midpoint from the preceding week. Older participants and the rest of young participants (N=96 young individuals) first completed an in-laboratory adaptation night to get familiar with the lab environment and allow for detection of any sleep disturbances. For the seven days before the baseline night, participants adhered to a regular sleep-wake schedule (within ±30 minutes) according to their habitual bed and wake-up times, verified using sleep diaries and wrist actigraphy (Actiwatch©, Cambridge Neurotechnology, UK). All participants were instructed to avoid daytime naps and to refrain from unusually intense physical activity during the final three days of the fixed sleep schedule. Their habitual sleep was then recorded in complete darkness using EEG. The current study focuses exclusively on the baseline night of sleep in both younger and late middle-aged individuals.

### EEG acquisitions

Polysomnographic sleep data of young participants were acquired using either V-Amp 16 amplifiers (Brain Products, Germany) (Ly et al., 2016; Muto et al., 2016, 2021; Vandewalle et al., 2009), a QuickAmp-72 (Brain Products, Germany) (Mascetti et al., 2013) or a N7000 amplifiers (EMBLA, Natus Medical Incorporated, Planegg, Germany) (Gaggioni et al., 2019). The electrode montage consisted of at least 9 EEG channels (F3, Fz, F4, C3, Cz, C4, Pz, O1, O2; reference to right mastoid), 2 bipolar EOGs, 2 bipolar EMGs, and 2 bipolar ECGs. For late-midlife group, participants sleep EEG data were acquired using N7000 amplifiers (EMBLA, Natus Medical Incorporated, Planegg, Germany) with 10 EEG channels (F3, Fz, F4, C3, Cz, C4, P3, Pz, P4, O1, O2), 2 bipolar EOGs, 2 bipolar EMGs, and 2 bipolar ECGs (Chylinski et al., 2021)]. For both age groups, acquisition on the screening night of sleep also included respiration belts, oximeter and nasal flow, 2 electrodes on one leg, but included only Fz, C3, Cz, Pz, Oz, and A1 channels. Baseline EEG data were re-referenced off-line to average mastoids.

### Arousal detection

Sleep stage scoring and arousal detection were carried out in separate steps by 2 independent algorithms. Sleep stage scoring was performed in 30-second windows using a validated algorithm (ASEEGA, Physip) (Berthomier et al., 2007; Peter-Derex et al., 2021). Automatic arousal detection was then computed as it is objective and reproducible (Chylinski et al., 2020). We used an individually tailored validated algorithm based on the AASM definition (Berry et al., 2012) of arousal but without using sleep stage information. Automatic scorings were visually inspected following computation.

In brief, arousal detection is performed over all electrodes on whole-night recordings split into 1-second epochs in 2 successive steps computed over the power in the broad-α (7–13 Hz), β (16–30 Hz), and lower-θ (3–7 Hz) frequency bands, excluding the σ band (11–16 Hz) — i.e., corresponding to frequency of sleep spindles — which cannot be considered as arousals. A fixed threshold is first applied to detect abnormal EEG activity relatively to the whole-night recording: any 1-second epoch with power in any of the 3 frequency bands higher than the whole-night median value in each frequency band is considered as a potential arousal. The second step adapts the threshold to account for the specific EEG background activity in a shorter time window. A specific threshold is computed for each 30-second window: all 1-second epochs without concomitant EMG tone increase are selected, as well as the first ten 1-second epochs without EMG increase before and after the 30-second window being evaluated; threshold of each frequency band consists in the median power over the selected 1-second epochs. Events composed of at least 3 consecutive 1-second epochs with changes in EEG frequencies higher than twice the local median and 1 median of the whole recording for that frequency band were considered as arousals. For detailed explanations on the method, see ref. (Chylinski et al., 2020).

As previously (Chylinski et al., 2021), we split arousal according to 2 criteria, which we considered as relevant in research settings, as well as clinical practice. The first criterion addressed whether arousals were associated with a sleep stage transition (T+) (when they occurred within 15 seconds of a stage change — in the second half of an epoch preceding a stage change or in the first half of an epoch assigned a different stage than the previous epoch) or whether they did not (T–). The second criterion considered the concomitant increase in EMG tone (M+) or its absence (M–).

We used the absolute number of arousals rather than arousal density to capture the total arousal burden across the night, which was our primary variable of interest. Normalizing to total sleep time (TST) can obscure meaningful differences when TST varies, specially between young and late middle-aged individuals. By including TST as a covariate, we were able to assess the independent contribution of arousal events to genetic risk for AD and cognitive outcomes, focusing on whether a greater overall arousal load—regardless of sleep duration—was linked to neurodegenerative vulnerability.

### Genotyping, quality control and imputation

Blood samples and buccal swabs were collected and stored at −20°C within few hours until DNA extraction. The genotyping was performed using the Illumina Infinium OmniExpress-24 BeadChip arrays (Illumina, San Diego, CA) based on Human Build 37 (GRCh37). All the study participants were European ancestry. Established quality control (QC) procedure was performed using PLINK (Purcell et al., 2007) (http://zzz.bwh.harvard.edu/plink/). In brief, the SNPs were excluded as follows: >10% missing genotypes, <95% call rate, minor allele frequency (MAF) below 0.01, out of Hardy-Weinberg equilibrium (p-value <10-4 for the Hardy-Weinberg test). SNPs on 23rd chromosome as well as ambiguous SNPs (A-T, T-A, C-G, G-C) were excluded as well. The data was matched for deviation with European ancestry using 1000 Genomes Project dataset (1KGP, https://www.internationalgenome.org). Imputation was conducted using the Sanger imputation server (https://imputation.sanger.ac.uk/) based on the Haplotype Reference Consortium (r1.1) as reference panel and using Eagle2.4 pre-phasing algorithm. The detailed data processing and analysis for young and late middle-aged sub-sample is as described previously in (Koshmanova et al., 2022; Muto et al., 2021). We finally ended with 7,165,614 SNPs common to all participants.

### Polygenic risk score (PRS) calculation

PRS analyses can be used to assess the genetic liability of an individual for a phenotype by calculating the weighted sum of risk alleles effect size identified in genome-wide association studies. In the current study, we calculated a PRS for AD for each participant using summary statistics from the recent GWAS meta-analysis of European ancestry (Wightman et al., 2021). This approach differed from the PRS calculation originally applied in part of the young individuals’ dataset (Muto et al., 2021). The standardization and quality control of GWAS summary statistics was performed by MungeSumstats, a Bioconductor R package (Murphy et al., 2021). In the process, the summary statistics was pruned to align reference alleles to build GRCh37, remove multiallelic variants, and adjust weights for the appropriate reference alleles. The PRS was then generated using SBayesR algorithm implemented in GCTB software. The approach assumes that the SNP effects are drawn from mixtures of distributions with the key metrics defining these genetic architectures estimated through Bayesian frameworks. To derive PRSs from GWAS effect estimates of SNPs, SBayesR essentially uses Bayesian linear mixed model and the reference linkage disequilibrium (LD) correlation matrix. In our analysis, we used banded LD matrix to improve the prediction accuracy as recommended by the authors of GCTB. We used p-value thresholding after PRS computation through PLINK to include only the SNPs reaching stringent GWAS significance (p-value <1×10^−8^) to restrict the number of genetic markers to a minimum.

### MRI data

MRI data were used in late middle-aged individuals in order to determine the region of interest used for extraction of Aβ burden value based on PET images, as fully described in (Chylinski et al., 2021). Quantitative multiparametric MRI acquisition was performed on a 3-Tesla MR scanner (Siemens MAGNETOM Prisma, Siemens Healthineers) to get a magnetization transfer–weighted (MT-weighted) contrast, based on multi-echo 3D fast low angle shot at 1 mm isotropic resolution (Weiskopf & Helms, 2008). MRI multiparameter maps were processed with the hMRI toolbox (Tabelow et al., 2019) (http://hmri.info) and SPM12 (Welcome Trust Centre for Neuroimaging, London, United Kingdom) to obtain a quantitative MT map and segmented images (gray matter, white matter, CSF), normalized to the standard MNI space using unified segmentation (Ashburner & Friston, 2005).

### PET scan

Aβ PET imaging was performed only in late middle-aged individuals as fully described in (Chylinski et al., 2021). Younger individuals typically have no detectable Aβ accumulation such that it would be unethical to expose them to radiation. Aβ PET imaging was performed using [^18^F]Flutemetamol, except for 3 volunteers for which [^18^F]Florbetapir was used as fully described in (Chylinski et al., 2021). PET scans were performed on an ECAT EXACT+ HR scanner (Siemens). Individual PET average images were manually reoriented according to MT-weighted structural MRI volumes and coregistered to the individual space structural MT map. Flow-field deformation parameters obtained from DARTEL spatial normalization of the MT maps were applied to averaged coregistered PET images (Ashburner, 2007). We did not provide correction for partial volume effect, as this type of PET processing was not included in Centiloid scaling pipeline (Klunk et al., 2015). Volumes of interest were determined using the automated anatomical labeling (AAL) atlas (Tzourio-Mazoyer et al., 2002). Standardized uptake value ratio (SUVR) was computed using the whole cerebellum as a reference region (Klunk et al., 2015). As images were acquired using 2 different radioligands, their SUVR values were converted into Centiloid units (Klunk et al., 2015). Aβ burden was averaged over a composite mask covering the previously reported earliest aggregation sites for Aβ pathology (Grothe et al., 2017) — frontal medial cortex and basal part of temporal lobe (fusiform and inferior temporal gyri). According to the study by Krasny et al. (2024), the amyloid PET positivity threshold was set at Centiloid 20.

### Cognitive assessment

Cognitive assessments were conducted at baseline in part of the younger sample (N=357, as part of the same unique study) while they were administered at baseline and at 2y (mean 767±54 days) and 7y (mean 2647±98 days) follow-up in the late middle-aged sample. Since significant associations between arousals and PRS for AD were observed only in the late middle-aged individuals (see Results), the following description focuses on the cognitive data from this group (the tests used for the young individuals were slightly different - for details see **Suppl. Fig. S7**). At the first time point, a cognitive battery of neuropsychological tasks was carried out in 2 sessions, while well rested. A first session of ~1 hour was performed in the afternoon prior to the sleep assessment, approximately 7.5 hours before habitual bedtime, and a second session of ~1.5 hours was performed on another day (between 12 and 6 hours prior to habitual bedtime). From those 2 sessions, 3 domain-specific composites scores were computed for the memory, executive function, and attentional domains, and they consisted of the standardized (z- scores) sum of the standardized domain-specific scores, where higher values indicate better performance.

The first session comprised (a) mnemonic similarity task (MST) (Stark et al., 2013); (b) category verbal fluency (letter and animals) (Cardebat et al., 1990); (c) digit symbol substitution task (DSST) (Tulsky et al., 1997); and (d) visual N-back task (1and 3-back variants) (Kirchner, 1958). The second session of ~1.5 hours was performed on another day (between 12 and 6 hours prior to habitual bedtime) and comprised (a) inverse order digit span task (Tulsky et al., 1997); (b) free and cued selective reminding test (FCSRT) (Grober et al., 1988); (c) trail making test (TMT) (Bowie & Harvey, 2006), and (d) and logical memory from Wechsler memory test (MEM-III) (Wechsler, 2001). The memory score consisted of the FCSRT (sum of all 4 free recalls), the recognition memory score from the MST, and delayed recall from the logical memory of MEM-III test. The executive function score included verbal fluency tests (letter and animals score for 2 minutes), inverse order digit span, TMT (part B minus part A), and N-back (3-back variant). The attentional score comprised the DSST, TMT (part A), and N-back (1-back variant).

The second assessment took place two years after the first, and the third was conducted seven years after the initial assessment. At these two time points, the same procedure and tests were used to compute the three composite scores. However, all individuals did not participate in the follow-up assessments and only 66 old individuals participated in the second time point assessment and 64 old individuals participated in the third time point assessment (with 48 individuals common to both follow-ups). The substantial dropout observed at the second cognitive assessment was primarily due to the COVID-19 crisis. For each composite score, cognitive decline was computed as the baseline performance minus the follow-up performance, divided by the baseline performance, so that a higher score indicates a higher decline over time.

### Statistics

Statistical analyses were performed in the R environment (version 4.1.3) (R Development Core Team, 2017) using generalized additive models for location scale and shape (GAMLSS) (Rigby & Stasinopoulos, 2005; D. M. Stasinopoulos & Rigby, 2008) based on the distribution of the dependent variables. GAMLSS offer a wide variety of family of distributions for model fitting (Rigby et al., 2019; Rigby & Stasinopoulos, 2014; M. D. Stasinopoulos et al., 2017) and provides more flexibility than GLM or GAM approaches. Outliers among all the assessed metrics lying beyond four times the standard deviation were removed from the analysis (the final number of individuals included in each analysis is reported in each table). Our primary objective was assessed in the first GAMLSS including the total number of arousals as the dependent variable and a four-way interaction among PRS for AD, transition status, EMG status, and age group as the main predictor. Since it included all variables of interest in the same model, it was not corrected for multiple comparisons and significance was set at p<0.05. Additionally, all related three-way and two-way interaction terms were included. Sex and total sleep time (TST) were added as covariates in the model and subjects were treated as random factors. This model yielded significant association between the four-way interaction and total number of arousals (see Results). Hence, a subsequent post-hoc GAMLSS analysis was used to specify which type of arousal was associated with the PRS for AD and in which age group this association was observed. This second GAMLSS included the PRS for AD as the dependent variable and four two-way interactions between each type of arousal (T+M+, T+M−, T−M+, T−M− arousal) and age group as independent variables. Sex and TST were added as covariates.

The association between arousal and cognition in each time point was assessed in models including each composite score as a dependent variable together with two types of arousals related to the PRS for AD (T+M+ and T+M− arousals) as independent variables as well as age, sex, education and TST as covariates (i.e. 9 models in total, 3 per time point). As an exploratory analysis, in similar models, we also considered the association between recognition memory score of MST in all three time points and T+M+ and T+M− arousals as this memory task is highly sensitive to early signs of cognitive decline (Marks et al., 2017; Stark et al., 2013).

We computed a priori sensitivity analysis to get an indication of the minimum detectable effect size in our main analyses given our sample size. According to G*Power 3 (version 3.1.9.4) (Faul et al., 2009), taking into account a power of .8, an error rate α of .05), a sample size of 540 allowed us to detect small effect sizes f^2^ >.023 (confidence interval: .008 –.063; R^2^> .022, R^2^ confidence interval: .008 –.059) within a linear multiple regression framework including four predictors (Arousals number, EMG status, transition status, age group) and 2 covariates (sex and TST).

We also computed a similar prior sensitivity analysis for the younger and late middle-aged groups separately, taking into account a power of .8, an error rate α of .05, a sample size of 453 young individuals allowed us to detect small effect sizes f^2^ =.032 (confidence interval: .02 –.09; R^2^ > .031, R^2^ confidence interval: .015 –.084); while a sample size of 87 late middle-aged adults allowed us to detect medium-large effect sizes f^2^ =.187 (confidence interval: .09 –.56; R^2^ > .157, R^2^ confidence interval: .082 –.358). There is limited published data on the precise effect sizes for quantitative sleep metrics and genetic associations in late middle-aged adults. However, the recent studies have used similar sample size in the analysis (Tsapanou et al., 2020) providing further support to the validity of our study results.

## Results

Comparing each type of arousals between the two age groups, we observe that the number of arousals associated with a sleep stage transition, i.e. T+M+ and T+M−, significantly increased in the late middle-aged individuals (**p<0.005**), while those not associated with a stage transition, i.e. T−M− & T−M+, decreased in this age group (**p<0.0001, Table 1**). Besides, age, body mass index (BMI), depression, sleep quality, and chronotype scores were significantly higher in the late middle-aged group compared to the younger group, whereas the younger group had a significantly longer total sleep time.

### Arousal heterogeneity reflects different associations with PRS for AD

Prior to seeking association between PRS and sleep metrics, we first assessed whether early PET Aβ burden and PRS for AD were correlated in late middle-aged individuals, since only this group underwent Aβ assessment. We did not find a significant association between Aβ levels and PRS for AD (**Figure 2A**; Sperman’s r=−.05; p=.64, and Sperman’s r=−.08; p=.43, when including the three Aβ-positive individuals - i.e. with centiloid values > 20- of the sample, **Suppl. Fig. S1**), supporting that PRS for AD is relatively independent of Aβ accumulation and grasps, at least in part, a distinct aspects of the AD-related risk as previously suggested by others (Ge et al., 2018; Luckett et al., 2022; Xicota et al., 2022).

**Figure 2.**
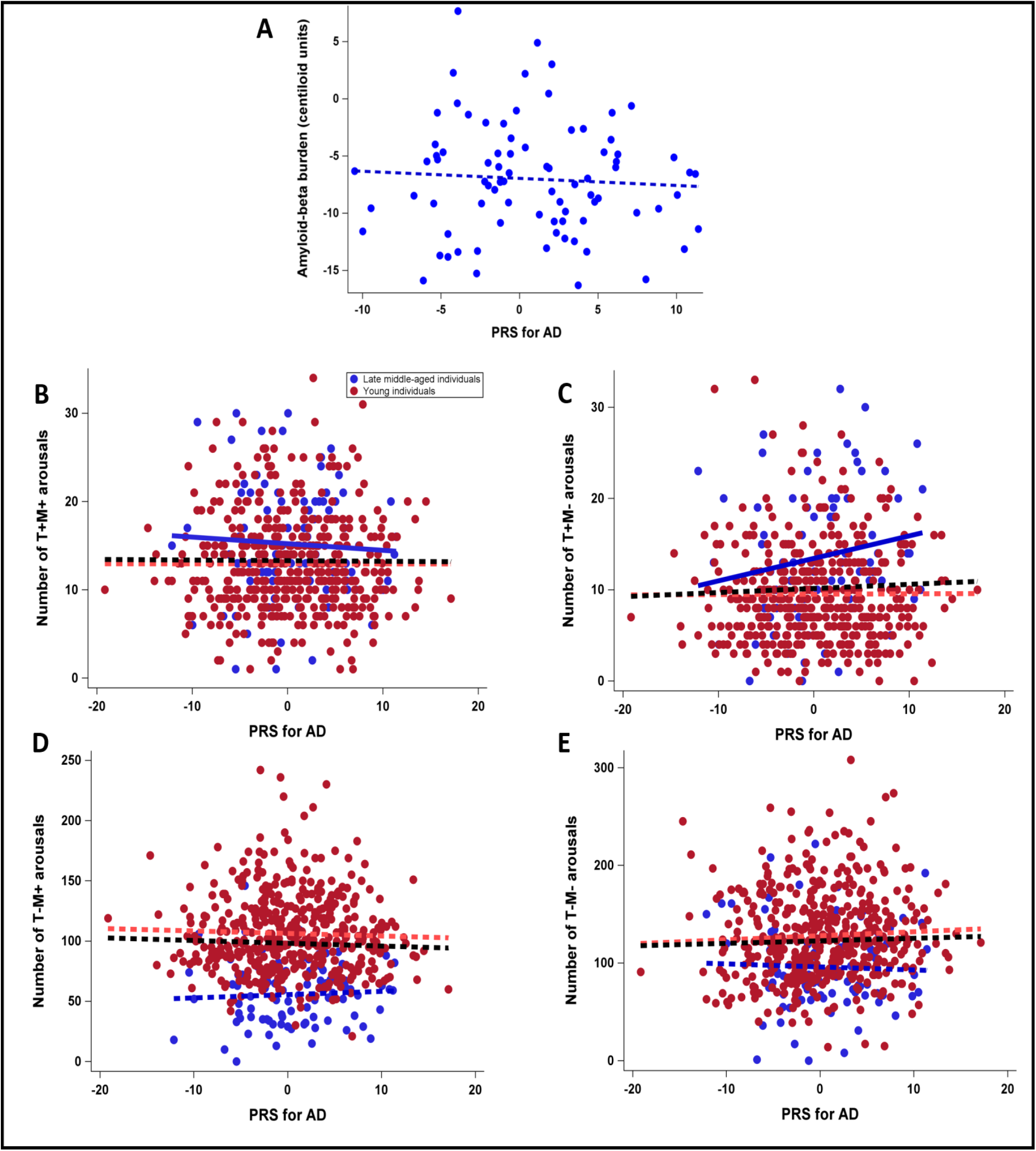
Associations between PRS for AD and sleep arousals. **(A)** Spearman correlation analysis showed that amyloid-beta burden and PRS for AD were not significantly correlated after excluding Aβ-positive individuals (N=82; p=0.64). Amyloid PET positivity was defined as a Centiloid value ≥ 20 (See Supplementary figure 1 for the association between PRS for AD and Aβ burden including Aβ-positive individuals). **(B)** Association between number of T+M+ arousals and the PRS for AD; age group by arousals interaction (N=540; p=0.05); post hoc subgroup analyses led to a significant negative association for the late middle-aged (p=0.01) but not the young group (p=0.83). **(C)** Association between number of T+M− arousals and the PRS for AD; age group by arousals interaction (N=536; p=.02); post hoc subgroup analyses led to a significant positive association for the late middle-aged (p=0.003) but not the young group (p=0.77). **(D)** Association between number of T−M+ arousals and the PRS for AD; age group by arousals interaction (p= N=539; 0.14). **(E)** Association between the number of T−M− arousals and the PRS for AD; age group by arousals interaction (N=540; p=0.13). Simple regression lines are used for a visual display and do not substitute the GAMLSS outputs (see Table 3). The black line represents the regression irrespective of age groups (young + old). Solid and dashed regression lines represent significant and non-significant outputs of the GAMLSS, respectively.

We then targeted our primary objective in a GAMLSS assessing whether arousal number, as dependent variable, was associated with PRS for AD across the different types of arousals (T+/T− & M+/M−), including age group, sex and total sleep time (TST) as covariates. The model yielded a significant 4-way interaction between PRS for AD, arousal transition and EMG statuses and age group (β=0.02; t= 2.52; **p=.01**), indicating that the association between arousals and PRS varied based on transition and EMG status as well as age group. Besides, the statistical model yielded significant main effects of transition status, age group, sex, TST, as well as significant two-way interactions between transition and EMG status, transition status and age group, EMG status and age group, and three-way interactions between PRS, and EMG and transition status, and between EMG and transition status and age groups (**Table 2**). Adding the variables that significantly differed between the two age groups (i.e., BMI, depression, sleep quality and chronotype) as a covariate to this primary GAMLSS did not change the statistical outputs although all these variables were significantly associated with number of arousals, indicating that the observed associations were unlikely to be driven by these group differences (**Suppl. Table S1**).

**Table 2.** Association between arousals number and PRS for AD (N=540). Analyses were performed using a generalized additive model for location, scale, and shape (GAMLSS). Results with p values less than 0.05 were considered statistically significant (df = 1801.64). Estimates in GAMLSS are reported in log scale. PRS: polygenic risk score; TST: total sleep time; EMG: Electromyography

| Independent variable | t | Estimate | SE | p-value |
| --- | --- | --- | --- | --- |
| PRS | -0.80 | -0.004 | 0.005 | 0.42 |
| Transition status | 41.43 | 1.29 | 0.03 | **<.0001** |
| EMG status | -1.00 | -0.04 | 0.04 | 0.32 |
| Age group | -7.15 | -0.29 | 0.03 | **<.0001** |
| Sex | 14.94 | 0.03 | 0.02 | **<.0001** |
| TST | 38.02 | 0.003 | 0.00006 | **<.0001** |
| PRS*Transition | 1.61 | 0.009 | 0.006 | 0.11 |
| PRS*EMG | 1.75 | 0.01 | 0.007 | 0.08 |
| Transition*EMG | 13.48 | 0.59 | 0.04 | **<.0001** |
| PRS* age group | 0.78 | 0.004 | 0.006 | 0.44 |
| Transition * age group | 23.91 | 0.80 | 0.03 | **<.0001** |
| EMG*age group | -5.83 | -0.31 | 0.04 | **<.0001** |
| PRS*Transition*EMG | -2.54 | -0.02 | 0.008 | **0.01** |
| PRS*Transition*age group | -1.73 | 0.01 | 0.006 | 0.08 |
| PRS*EMG*age group | -1.24 | -0.01 | 0.008 | 0.22 |
| Transition*EMG*age group | -2.27 | -0.08 | 0.05 | **0.02** |
| PRS*Transition*EMG*age group | 2.52 | 0.02 | 0.009 | **0.01** |

To understand what was driving the 4-way interaction, we computed a post hoc GAMLSS with PRS, as a dependent variable, and the interaction between the number of each arousal type and age group as covariate, including their interactions (4 two-way interactions) regressing out TST and sex. The model yielded a significant link between PRS and the interaction between T+M− arousals and age group (β=−0.32; t=−2.40; **p=.02**) as well as a nominally significant link with the interaction between T+M+ arousals and age group (β=0.29; t=1.96; **p=.05**) on top of the main effect of T+M+ and T+M− arousals number (**Figure 2B-C**, **Table 3**). The other associations, including the association between PRS and interaction between T− M− and T−M+ arousals and age group, were not significant (**Figure 2D-E**, **Table 3**). Post hoc contrasts showed that there is a negative association between PRS and T+M+ arousals (β=−.30; t=−2.57; **p=.01**) as well as a positive association between T+M− arousals (β=.29; t=3.09; **p=.003**) in the late middle-aged group but not the younger group (β=.01; t=.22; p=.83 and β=.02; t=.29; p=.77 respectively; **Figure 2B-C**). Hence, the post hoc analysis indicates that the significant 4-way interaction found in the primary analysis is driven by the association between PRS and both T+M+ arousals and T+M− arousals in the late middle-aged group.

**Table 3.** Association between PRS for AD and the interaction between each arousal type and age group (N=540). Analyses were performed using a generalized additive model for location, scale, and shape (GAMLSS). Results with p values less than 0.05 were considered statistically significant. Estimates in GAMLSS are reported in log scale. Prior to the analysis, we removed the outliers among all variables by excluding the samples lying beyond four times the standard deviation. TST: total sleep time.

| Independent variable | t | Estimate | SE | p-value |
| --- | --- | --- | --- | --- |
| T+M+ arousals number | -1.93 | -0.30 | 0.13 | **0.05** |
| T-M+ arousals number | 1.28 | 0.04 | 0.03 | 0.20 |
| T+M- arousals number | 2.49 | 0.30 | 0.11 | **0.01** |
| T-M- arousals number | -1.19 | -0.02 | 0.02 | 0.23 |
| T+M+ arousals number *age group | 1.96 | 0.29 | 0.15 | **0.05** |
| T-M+ arousals number *age group | -1.50 | -0.04 | 0.03 | 0.14 |
| T+M- arousals number *age group | -2.40 | -0.32 | 0.13 | **0.02** |
| T-M- arousals number *age group | 1.52 | 0.03 | 0.02 | 0.13 |
| group | -0.22 | -2.33 | 2.59 | 0.83 |
| sex | -0.11 | -1.23 | 1.45 | 0.91 |
| TST | -0.29 | 0.0009 | 0.007 | 0.77 |

Given the sex imbalance of our samples (particularly in the young sample), we further assessed sex differences and could not find evidence that they were contributing to our finding (**Suppl. Table S2; Suppl. Figure S2-S3**). Besides, we could not find indications that the association between PRS and T+M+ arousals in late middle-aged individuals was driven by arousals detected during REM or NREM sleep (according to ASSM criteria, arousals must be associated with muscle tone increase during REM, such that all M− arousal were only detected during NREM) but rather found that it is the total number of T+M+ in REM and NREM that was associated with the PRS for AD (**Suppl. Table S3**).

### Arousals linked with PRS for AD are associated with cognitive performance and cognitive decline in late middle-aged individuals

We then tested whether cognitive performance of the late middle-aged individuals was differentially associated with the 2 arousal types that showed opposite association with PRS for AD. The GAMLSSs for each time point included each cognitive domain as dependent variable (i.e. three GAMLSS at each time point) and T+M+ arousals and T+M− arousals as covariates together with sex, age, education and TST. Considering the baseline assessment, GAMLSSs yielded a negative association between attentional domain and T+M− arousals (β=−06; t=−2.26; **p=.02**) and a positive association between attentional domain and T+M+ arousals (β=.08; t=2.09; **p=.04**) on top of the effect of age and education (**Table 4**). There was no significant association between these arousals and the other cognitive domains (i.e. memory and executive functions; p>.10; **Table 4**, **Figure 3A-B, Suppl. Figure S4**). Adding PRS as a covariate to these models did not change the statistical outputs while PRS itself was not significantly associated with the cognitive performance (p>.93).

**Figure 3.**
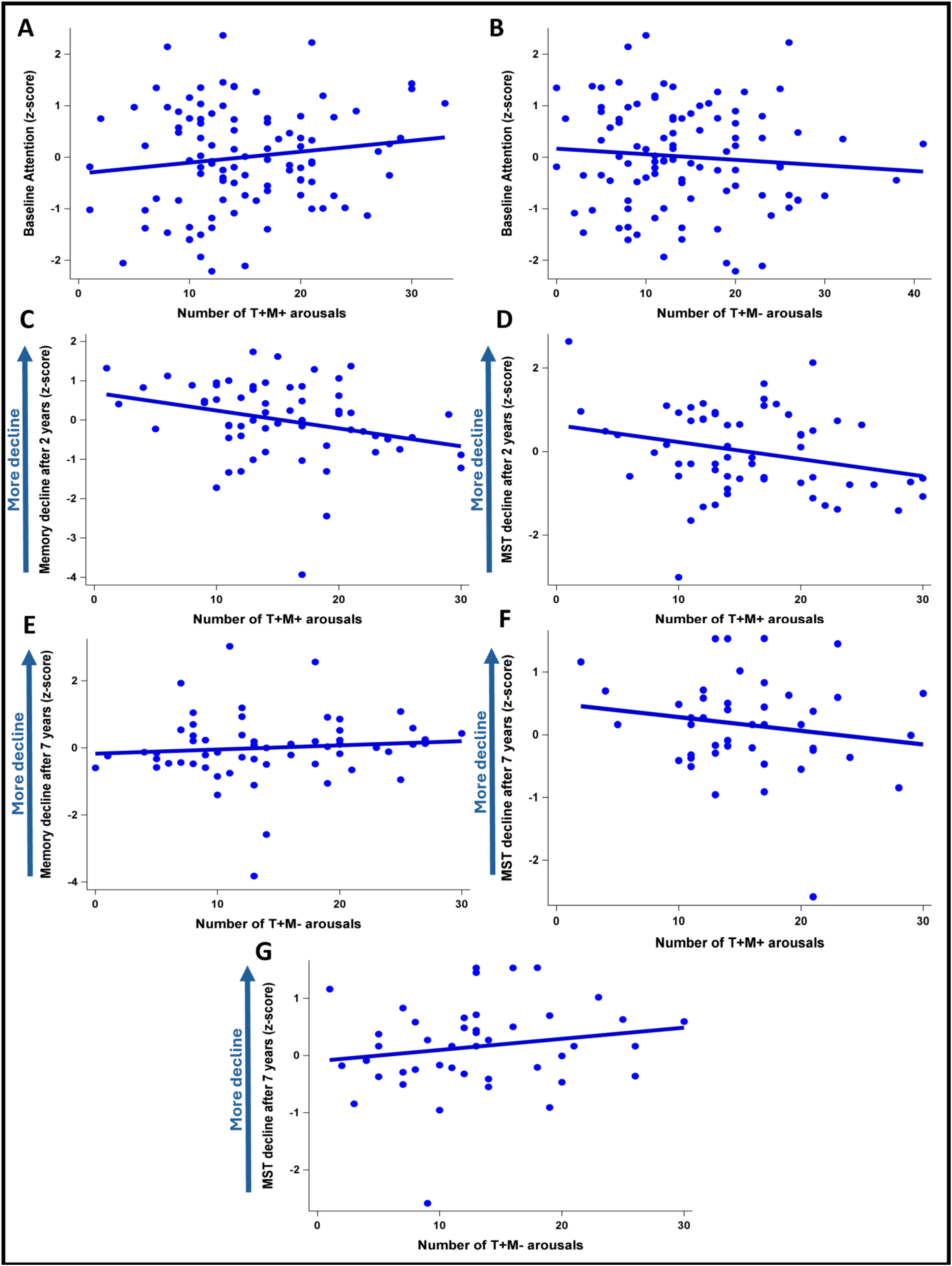
Significant associations between sleep arousals and cognitive scores. **(A)** Significant positive association between the number of T+M+ arousals and baseline attention scores (N=87; p=0.03). **(B)** Significant negative association between the number of T+M− arousals and baseline attention scores (N=87; p=0.02). **(C)** Significant negative association between the number of T+M+ arousals and memory decline after 2 years (N=61; p=0.002). **(D)** Significant negative association between the number of T+M+ arousals and the MST decline after 2 years (N=64; p=0.01). **(E)** Significant positive association between the number of T+M− arousals and memory decline after 7 years (N=60; p=0.02). **(F)** Significant negative association between the number of T+M+ arousals and MST decline after 7 years (N=47; p=0.006). **(G)** Significant association between the number of T+M− arousals and MST decline after 7 years. Although the Spearman correlation was not significant and the positive association cannot be observed in the plot, after controlling for age, sex, education and total sleep time, the GAMLSS yielded a significant positive association between the number of T+M− arousals and MST decline after 7 years (p=0.04). Simple regression lines are used for a visual display and do not substitute the GAMLSS outputs (Table 4). Solid regression line represent significant outputs of the GAMLSS. See Supplementary figure S2-4 for the non-significant associations between sleep arousals and cognitive scores.

**Table 4.**
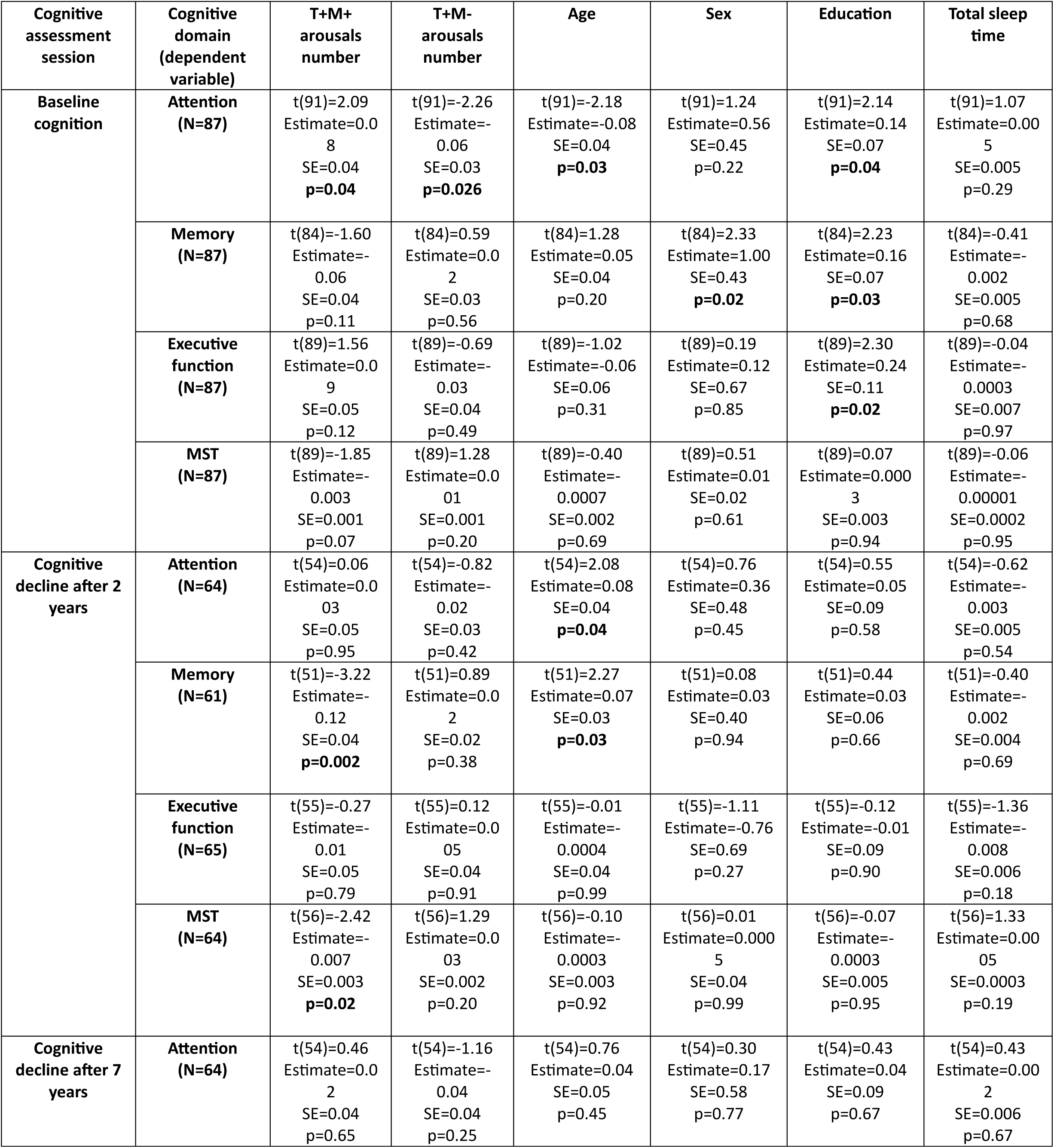
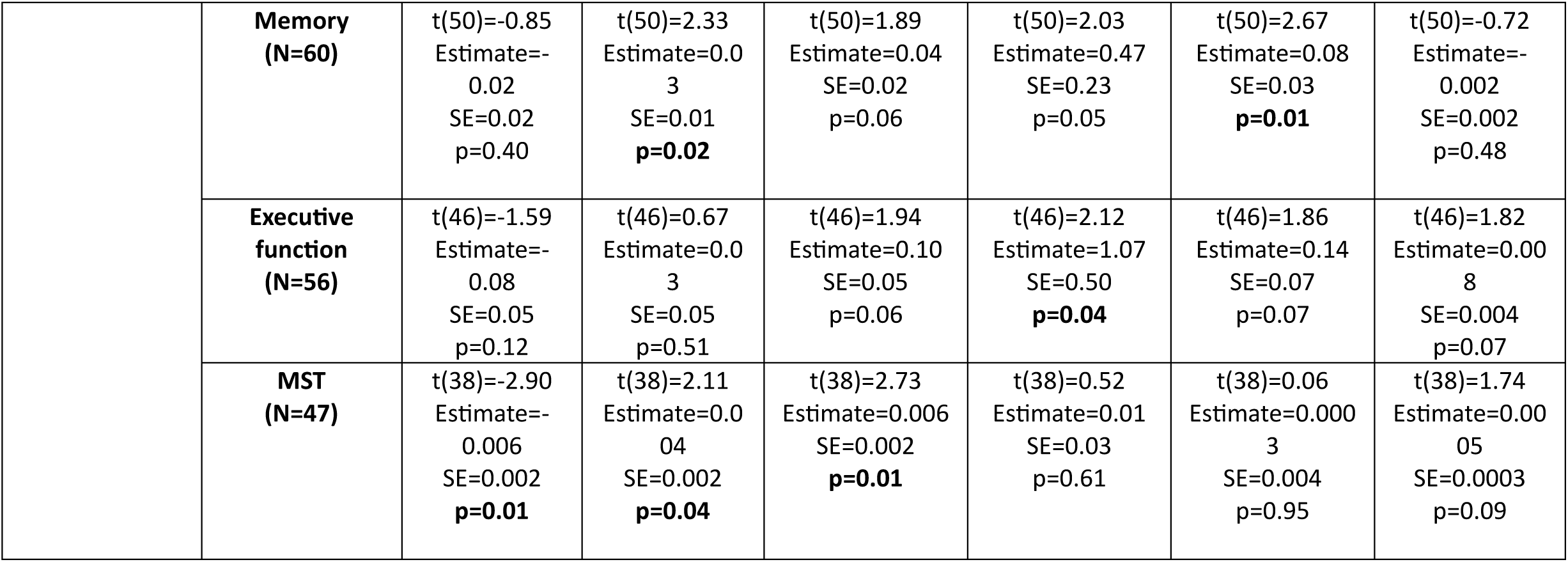
Association between cognitive domains at each time point and T+M+ and T+M− arousals in late middle-aged individuals. Analyses were performed using a generalized additive model for location, scale, and shape (GAMLSS). Results with p values less than 0.05 were considered statistically significant. Estimates in GAMLSS are reported in log scale. Prior to the analysis, we removed the outliers among all variables by excluding the samples lying beyond four times the standard deviation (the final number of individuals included in each analysis is reported below each dependent variable). MST: mnemonic similarity task.

We then focused on our first follow-up at 2 years that was completed in part of the late middle-aged individuals (N=66). The GAMLSSs led to a significant negative association between T+M+ arousals and performance change over the memory domain, indicating that more T+M+ arousals at baseline are associated with less memory decline over 2 years (β=−.12; t=−3.22; **p=.002**), on top of a main effect of age (**Figure 3C**, **Table 4**). The associations with the performance change in the other cognitive domains were not significant (p>.37; **Table 4**, **Suppl. Figure S5**). Adding PRS as a covariate to these models did not change the statistical output while PRS itself was not significant (p>.09).

We further considered cognitive decline after 7 years that was assessed in a subset of the late middle-aged participants (N=64; 73% overlap with 2y follow up – see methods). GAMLSSs yielded a positive association between memory decline and T+M− arousals (β=.03; t=2.33; **p=.02**), on top of the effect of education (**Table 4**, **Figure 3E**). Again, the associations with other cognitive domains were not significant (p>.37; **Table 4**, **Figure 3, Suppl. Figure S6**). Again also, adding PRS as a covariate to this model did not change the results while PRS itself was not significant (p>.16).

In a final step, we considered changes in the performance to the MST task, as it is a memory test reported to be highly sensitive to early signs of cognitive decline (Marks et al., 2017). We found a significant negative association between MST performance decline after two years and T+M+ arousals (β=−.007; t=−2.42; **p=.02; Figure 3D**, **Table 4**). Adding PRS as a covariate to this model did not change the results while PRS itself was not significant (p>.69). A subsequent GAMLSS also yielded a significant negative association between MST performance decline after seven years and T+M+ arousals (β=−.006; t=−2.90; **p=.01**) and a positive association between MST decline after seven years and T+M− arousals (β=.004; t=2.11**p=.04**) on top of a main effect of age (**Table 4**, **Figure 3F-G, Suppl. Figure S4, S5**). Again, adding PRS as a covariate to this model did not change the results while PRS itself was not significant (p>.70). Of note, none of the cognitive results survived false discovery rate (FDR) correction for the 18 tests performed (3 time points × 2 arousal metrics × 3 main cognitive domains), meaning these associations should be interpreted with caution. Finally, we found no significant association between T+M+ or T+M− and baseline performance in young individuals in any cognitive domain, suggesting the associations we uncovered previously are specific to the late middle-aged individuals of our sample (**Suppl. Table S4; Suppl. Figure S7**).

## Discussion

Sleep arousals are commonly viewed as disruptive and leading to negative functional outcomes (Mahowald & Schenck, 2005). This likely arises in part from their increased prevalence with aging, their association with external disturbances (e.g. noise) or with hypopnea/apnea in patients with sleep disordered breathing (SDB). Arousals are also triggered spontaneously, i.e. without a concomitant disturbance, and these may serve beneficial purposes for brain functioning (Halasz & Bodizs, 2012). In addition, spontaneous arousal are heterogenous, likely in part because of differential activity of the LC (Foustoukos & Lüthi, 2025; Osorio-Forero et al., 2025), and this heterogeneity is associated with early Aβ burden (Chylinski et al., 2021). In the present study, we determined whether spontaneous arousals are associated with the risk for developing AD and with cognition in healthy younger and late middle-aged individuals. We show that, depending on whether they are associated with a transition of sleep stage and with an increase in muscle tone, arousal can be associated with an increased or decreased polygenic risk for AD in late middle-aged, but not in younger adults. We further show that, in late middle-aged individuals, arousals heterogeneity is associated not only with current attentional performance but also bears predictive value for memory performance at 2 or 7 years. Although the effect sizes we observed were modest, if persistent, these small effects could influence lifelong health trajectories. These findings add to the growing literature showing that alterations in microstructural elements of sleep electrophysiology are associated with early AD neurobiology and precede the onset of AD symptoms (Chylinski et al., 2021, 2022; Ju et al., 2017; Sharon et al., 2025). They may contribute to establishing a marker of AD risk in otherwise healthy individuals.

We showed, in the same sample of late middle-aged individuals, that defining arousal based on stage transition and association with muscle tone was meaningful as arousal types differed in their oscillatory compositions, but – in absence of a younger group - could not report age related changes across the different types of arousals (Chylinski et al., 2021). Here, we report that, contrary to the general assumption, not all arousal types increase in the late middle-aged group. Only the number of arousals that are associated with sleep stage transition, and therefore most impinging on sleep continuity, increase with age, while those not associated with these transitions become less common. This constitutes the first important observation that arousal heterogeneity is meaningful. In addition, it is precisely those arousals increasing with age that we find associated with the PRS for AD. In other words, the extent of the age-related difference in T+ is linked to the genetic risk for developing AD.

We find that, in late middle-aged individuals, more T+M− arousals are associated with an increased PRS for AD, which is similar to the association between T+M− arousals and higher early amyloid burden we previously reported in the same sample (Chylinski et al., 2021). Arousals associated with sleep stage transition but not with a detectable muscle tone increase appear therefore to be related to more risk for developing AD both based on genetic and protein burden assessment. T+M− arousals may therefore reflect a chronic sleep disruption that contributes to, or is at least associated with, long-term neural vulnerability to AD-related processes. In line with this negative view, the amount of T+M− arousals is also associated with poorer attentional performance at baseline and memory decline at 7y (as shown using the global memory score and the exploration of the MST, a task sensitive to early hippocampal dysfunction (Marks et al., 2017), which supports the idea that attentional deficits may precede and potentially exacerbate memory deterioration over time. In contrast to T+M− arousals and still in the late middle-aged participants, we find that the amount of T+M+ arousals is associated with a reduced PRS for AD, better attentional performance at baseline and better memory performance at 2y and 7y (shown through both the global memory and MST scores at 2y and the MST score at 7y). Hence, these arousals would represent a better situation in terms of AD risk, associated with better attention processes which could favor subsequent better memory function.

According to our findings, the main feature differentiating negative and positive transition arousals is whether or not they are accompanied by an increase in EMG tone. Since by definition an arousal that would exclusively consist of a muscle activation (without EEG activation) do not exist, arousals with muscle tone increase were proposed to represent stronger brain activation (Halász et al., 2004). A study in mice showed that cortico-hippocampal coherence increases prior to and during arousals accompanied by EMG (dos Santos Lima et al., 2019). It is also shown that stronger arousals that are accompanied by increased muscle tone are associated with the appearance of theta activity in the hippocampus (Jarosiewicz & Skaggs, 2004), while theta activity of hippocampus has been reported to be important for memory processing (Boyce et al., 2016).

Although multiple brain regions contribute to the regulation of arousal - for example an EEG-fMRI study in humans showed that the midbrain, thalamus, basal ganglia, and cerebellum, were activated during arousal while cortical regions were deactivated (Zou et al., 2020), the LC may be among the most important effector. Studies in animals showed that optogenetic activation of the LC, in the midbrain, causes arousals associated with EMG both during REM and NREM sleep, most often leading to full wakefulness (Carter et al., 2010). More importantly, spontaneous arousals in rodents were recently reported to align with variations in noradrenaline (arising from LC activity) leading to two types of arousal (Foustoukos & Lüthi, 2025; Osorio-Forero et al., 2025). The LC appears to regulate arousal intensity, with most LC surges (70%) not leading to full wakefulness. Only stronger surges (30%) cause brief awakenings, while smaller ones induce partial arousal affecting the heart and thalamus but not the cortex (Osorio-Forero et al., 2025). The LC, and also potentially other subcortical structures, may underlie arousal heterogeneity, particularly those associated with EMG changes. According to our results, these may be the most profitable, whether or not they are strong enough to lead to a sleep stage transition (sleep stage being a somewhat arbitrary classification). In that respect T+M− arousal may represent incomplete arousal events in which the full arousal network is not recruited or arousal arising from a distinct set of brain regions and, according to our findings, would be associated with more deleterious outcomes.

We emphasize that in our previous work on arousal heterogeneity, we found that T−M+ arousals were associated with reduced Aβ burden and better attention performance (Chylinski et al., 2021), which is only partially in line with our current findings, where T+M+ are associated with lower PRS for AD and better attention. We may have lacked sensitivity to find positive associations with both T+M+ and T−M+ arousals across both analyses. The discrepancy may also arise from the fact that we used two different approaches that do not grasp exactly the same part of the risk for AD. Consistent with previous research (Leonenko et al., 2019), we indeed found no association between PRS for AD and Aβ burden. PRS may capture a broader genetic risk for AD and may be more predictive of disease progression than amyloid accumulation itself (Leonenko et al., 2019). In our earlier study, we interpreted the sleep-stage transition component as the key feature linking arousals to AD risk. Taken together, however, the two studies may suggest the following: while T– arousals may be associated with lower Aβ accumulation and T+ arousals with reduced genetic vulnerability, arousals accompanied by EMG activation—regardless of stage transition—consistently mark lower risk for future AD. This pattern supports the idea that motor-tone recruitment during arousals reflects a more complete and coordinated activation of the arousal system, which may have protective or compensatory value in maintaining cognitive function. One could speculate that M+ arousals offer recurring opportunities to transiently synchronize distant brain areas, similarly to sleep spindles (Steriade, 2003). This warrant future investigation in a distinct, and ideally larger, sample of late middle-aged or older individuals.

Despite having a much larger sample, we could not isolate associations between arousal types and PRS in young adults. This could mean that it is only when brain function and sleep become more susceptible to challenges (Schmidt et al., 2012) that association between arousal and the biology of AD emerges in those that are more prone to develop AD. Sleep arousals would in turns not constitute a promising marker of AD risk in young adults. Yet in late middle-aged individuals, (T+M− and T+M+) arousals, and not the PRS for AD, were associated with cognitive changes over 2 or 7 years. Sleep microstructure could therefore constitute a promising early marker of future cognition and brain ageing trajectory that is more sensitive that risk assessment based on both genetics and amyloid beta burden assessments (Chylinski et al., 2021).

We acknowledge several limitations of our study. First, the number of participants who completed the follow-up cognitive assessment was relatively small (N= 66 and N = 64) and the number of participants that completed both follow-up assessment was even smaller (N = 48 – see methods). Our sensitivity at follow-ups was therefore lower and may have hindered other weaker significant associations such that our results have to be taken mostly in relative terms (stronger vs. weaker effect rather than presence vs. absence of effects). In addition, the memory composite and the MST scores were not consistently associated with T+M+ and T+M− arousals across the 2 follow-up assessments at 2 and 7y. It is rather the overall pattern of associations across both scores that is consistent with memory performance changes being associated with arousal types. Whether this precise pattern reflects a lack of sensitivity or actual differences over different aspects of memory function is not known. Besides, the exclusion criteria applied were stringent and do not reflect the variability present in the general population. Although this guarantees that our findings are not biased by common age-related health issues, our participants exhibited minimal Aβ accumulation, with few individuals classified as Aβ-positive (N=3). In particular, since we did not included patients with SDB, our findings likely do not generalize to the perturbation-induced arousals, which have been associated with adverse behavioral (Aloia et al., 2004; Roehrs et al., 1994) and neurodegenerative outcomes (Ju et al., 2017). Furthermore, the predictive validity of PRS remains a subject of ongoing debate (Koch et al., 2023) and we cannot determine which participants will ultimately develop AD. Moreover, as already stated and even though it was controlled for, sex was not balanced within and across age groups while women exhibit distinct sleep characteristics (Eggert et al., 2021; Li et al., 2022), and are also more susceptible to AD (Andrew & Tierney, 2018). All these aspects limit the generalizability of our finding and warrants future investigations including larger and sex-balanced samples with more lenient exclusion criteria, e.g. including OSD and additional / more detailed memory performance assessments.

## Conclusion

By leveraging EEG data and genotyping across a diverse age range, we provide novel insights into how subtle features of sleep architecture may reflect or interact with underlying genetic susceptibilities. We report that spontaneous arousals during sleep are not uniform but vary in their association with genetic risk for AD and bears prediction value for future cognitive decline depending on their electrophysiological characteristics. These associations emerged only in older individuals, consistent with age-related vulnerability to subtle sleep disruptions. By distinguishing arousal subtypes, our study highlights sleep microstructure as a potential early marker of neurodegenerative risk during early aging. How arousal may interact with other features of sleep microstructure to contribute to AD risk remains to be assessed.

## Author contribution

Conceptualization, N.M., G.V. and P.T.; Methodology, N.M., P.T., M.Z.; Software, N.M.; Validation, G.V., F.C., and C.B.; Formal Analysis, N.M., P.T.; Investigation, N.M., G.V., P.T.; Resources, F.C., C.B., P.M., and G.V.; Data Curation, N.M., P.T.; Writing – Original Draft Preparation, N.M., G.V. and P.T.; Writing – Review & Editing, all authors; Visualization, N.M.; Supervision, G.V.; Project Administration, N.M. and G.V.; Funding Acquisition, F.C., C.B., P.M., and G.V. All authors have read and agreed to the published version of the manuscript.

## Acknowledgement

PT is supported by the EU Joint Programme Neurodegenerative Disease Research (JPND) IRONSLEEP project (FNRS reference: PINT-MULTI R.8011.21). FC and CB and GV are supported by the Fonds de la Recherche Scientifique - FNRS-Belgium. The study was supported by the Wallonia-Brussels Federation (Actions de Recherche Concertées - ARC—09/14-03), WELBIO/Walloon Excellence in Life Sciences and Biotechnology Grant (WELBIOCR-2010-06E), FNRS-Belgium (FRS-FNRS, F.4513.17 and T.0242.19 and 3.4516.11), Fondation Recherche Alzheimer (SAO-FRA 2019/0025), University of Liege (ULiege), Fondation Simone et Pierre Clerdent, European Regional Development Fund (Radiomed project), Fonds Leon Fredericq, EU JPND program (IRONSLEEP project - PINT-MULTI R.8011.21). The authors also thank Sarah Chellappa, Annick Claes, Catherine Hagelstein, Gregory Hammad, Brigitte Herbillon, Patrick Hawotte, Mathieu Jaspar, Sophie Laloux, Erik Lambot, Benjamin Lauricella, André Luxen, Julien Ly, Christelle Meyer, and Eric Salmon for their help over the different steps of the study. This work was conducted at the GIGA-In Vivo Imaging platform of ULiège, Belgium. The genotyping was performed at Genomics platform of GIGA institute at ULiège, Belgium.

## Data availability

The data and analysis scripts supporting the results included in this manuscript are publicly available via https://gitlab.uliege.be/CyclotronResearchCentre/Public/xxx (to be done following peer reviewing and upon acceptance for publication/and editor request). Researchers willing to access the raw data should send a request to the corresponding author (GV). Data sharing will require evaluation of the request by the local Research Ethics Board and the signature of a data transfer agreement (DTA).

## Conflicts of Interest

The authors declare no conflict of interest.

## Notes

### Competing Interest Statement

The authors have declared no competing interest.

## References

Aloia, M. S., Arnedt, J. T., Davis, J. D., Riggs, R. L., & Byrd, D. (2004). Neuropsychological sequelae of obstructive sleep apnea-hypopnea syndrome: a critical review. Journal of the International Neuropsychological Society, 10(5), 772–785.

Andrew, M. K., & Tierney, M. C. (2018). The puzzle of sex, gender and Alzheimer’s disease: Why are women more often affected than men? Women’s Health, 14, 1745506518817995.

Ashburner, J. (2007). A fast diffeomorphic image registration algorithm. Neuroimage, 38(1), 95–113.

Ashburner, J., & Friston, K. J. (2005). Unified segmentation. Neuroimage, 26(3), 839–851.

Baker, E., & Escott-Price, V. (2020). Polygenic risk scores in Alzheimer’s disease: current applications and future directions. Frontiers in Digital Health, 2, 14.

Beck, A. T., Epstein, N., Brown, G., & Steer, R. A. (1988). An inventory for measuring clinical anxiety: psychometric properties. Journal of Consulting and Clinical Psychology, 56(6), 893.

Beck, A. T., Steer, R. A., & Carbin, M. G. (1988). Psychometric properties of the Beck Depression Inventory: Twenty-five years of evaluation. Clinical Psychology Review, 8(1), 77–100.

Berry, R. B., Brooks, R., Gamaldo, C. E., Harding, S. M., Marcus, C., & Vaughn, B. V. (2012). The AASM manual for the scoring of sleep and associated events. *Rules, Terminology and Technical Specifications, Darien, Illinois*, American Academy of Sleep Medicine, 176, 2012.

Berthomier, C., Drouot, X., Herman-Stoïca, M., Berthomier, P., Prado, J., Bokar-Thire, D., Benoit, O., Mattout, J., & D’Ortho, M.-P. (2007). Automatic analysis of single-channel sleep EEG: validation in healthy individuals. Sleep, 30(11), 1587–1595.

Bowie, C. R., & Harvey, P. D. (2006). Administration and interpretation of the Trail Making Test. Nature Protocols, 1(5), 2277–2281.

Boyce, R., Glasgow, S. D., Williams, S., & Adamantidis, A. (2016). Causal evidence for the role of REM sleep theta rhythm in contextual memory consolidation. Science, 352(6287), 812–816.

Braak, H., & Del Tredici, K. (2012). Where, when, and in what form does sporadic Alzheimer’s disease begin? Current Opinion in Neurology, 25(6), 708–714.

Buysse, D. J., Reynolds III, C. F., Monk, T. H., Berman, S. R., & Kupfer, D. J. (1989). The Pittsburgh Sleep Quality Index: a new instrument for psychiatric practice and research. Psychiatry Research, 28(2), 193–213.

Cardebat, D., Doyon, B., Puel, M., Goulet, P., & Joanette, Y. (1990). Formal and semantic lexical evocation in normal subjects. Performance and dynamics of production as a function of sex, age and educational level. Acta Neurologica Belgica, 90(4), 207–217.

Carter, M. E., Yizhar, O., Chikahisa, S., Nguyen, H., Adamantidis, A., Nishino, S., Deisseroth, K., & De Lecea, L. (2010). Tuning arousal with optogenetic modulation of locus coeruleus neurons. Nature Neuroscience, 13(12), 1526–1533.

Chylinski, D., Rudzik, F., Coppieters ‘t Wallant, D., Grignard, M., Vandeleene, N., Van Egroo, M., Thiesse, L., Solbach, S., Maquet, P., & Phillips, C. (2020). Validation of an automatic arousal detection algorithm for whole-night sleep EEG recordings. Clocks & Sleep, 2(3), 258–272.

Chylinski, D., van Egroo, M., Narbutas, J., Grignard, M., Koshmanova, E., Berthomier, C., Berthomier, P., Brandewinder, M., Salmon, E., Bahri, M. A., Bastin, C., Collette, F., Phillips, C., Maquet, P., Muto, V., & Vandewalle, G. (2021). Heterogeneity in the links between sleep arousals, amyloid-β, and cognition. JCI Insight, 6(24). 10.1172/jci.insight.152858

Chylinski, D., Van Egroo, M., Narbutas, J., Muto, V., Bahri, M. A., Berthomier, C., Salmon, E., Bastin, C., Phillips, C., & Collette, F. (2022). Timely coupling of sleep spindles and slow waves is linked to early amyloid-β burden and predicts memory decline. ELife, 11, e78191.

dos Santos Lima, G. Z., Lobao-Soares, B., Corso, G., Belchior, H., Lopes, S. R., de Lima Prado, T., Nascimento, G., França, A. C. de, Fontenele-Araújo, J., & Ivanov, P. C. (2019). Hippocampal and cortical communication around micro-arousals in slow-wave sleep. Scientific Reports, 9(1), 5876.

Eggert, T., Dorn, H., & Danker-Hopfe, H. (2021). Nocturnal brain activity differs with age and sex: comparisons of sleep EEG power spectra between young and elderly men, and between 60–80-year-old men and women. Nature and Science of Sleep, 1611–1630.

Escott-Price, V., Sims, R., Bannister, C., Harold, D., Vronskaya, M., Majounie, E., Badarinarayan, N., Morgan, K., Passmore, P., Holmes, C., Powell, J., Brayne, C., Gill, M., Mead, S., Goate, A., Cruchaga, C., Lambert, J. C., Van Duijn, C., Maier, W., … Williams, J. (2015). Common polygenic variation enhances risk prediction for Alzheimer’s disease. Brain, 138(12), 3673–3684. 10.1093/brain/awv268

Escott-Price, V., Myers, A. J., Huentelman, M., & Hardy, J. (2017). Polygenic risk score analysis of pathologically confirmed Alzheimer disease. Annals of Neurology, 82(2), 311–314.

Faul, F., Erdfelder, E., Buchner, A., & Lang, A.-G. (2009). Statistical power analyses using G* Power 3.1: Tests for correlation and regression analyses. Behavior Research Methods, 41(4), 1149–1160.

Fleisher, A. S., Chen, K., Liu, X., Ayutyanont, N., Roontiva, A., Thiyyagura, P., Protas, H., Joshi, A. D., Sabbagh, M., & Sadowsky, C. H. (2013). Apolipoprotein E ε4 and age effects on florbetapir positron emission tomography in healthy aging and Alzheimer disease. Neurobiology of Aging, 34(1), 1–12.

Foustoukos, G., & Lüthi, A. (2025). Monoaminergic signaling during mammalian NREM sleep-Recent insights and next-level questions. Current Opinion in Neurobiology, 92, 103025.

Gaggioni, G., Ly, J. Q. M., Muto, V., Chellappa, S. L., Jaspar, M., Meyer, C., Delfosse, T., Vanvinckenroye, A., Dumont, R., & Coppieters’t Wallant, D. (2019). Age-related decrease in cortical excitability circadian variations during sleep loss and its links with cognition. Neurobiology of Aging, 78, 52–63.

Ge, T., Sabuncu, M. R., Smoller, J. W., Sperling, R. A., Mormino, E. C., & Initiative, A. D. N. (2018). Dissociable influences of APOE ε4 and polygenic risk of AD dementia on amyloid and cognition. Neurology, 90(18), e1605–e1612.

Grober, E., Buschke, H., Crystal, H., Bang, S., & Dresner, R. (1988). Screening for dementia by memory testing. Neurology, 38(6), 900.

Grothe, M. J., Barthel, H., Sepulcre, J., Dyrba, M., Sabri, O., & Teipel, S. J. (2017). In vivo staging of regional amyloid deposition. Neurology, 89(20), 2031–2038. 10.1212/WNL.0000000000004643

Halasz, P., & Bodizs, R. (2012). Dynamic structure of NREM sleep. Springer Science & Business Media.

Halász, P., Terzano, M., Parrino, L., & Bódizs, R. (2004). The nature of arousal in sleep. Journal of Sleep Research, 13(1), 1–23.

Horne, J. A., & Ostberg, O. (1976). A self-assessment questionnaire to determine morningness-eveningness in human circadian rhythms. International Journal of Chronobiology, 4(2), 97–110.

Jarosiewicz, B., & Skaggs, W. E. (2004). Level of arousal during the small irregular activity state in the rat hippocampal EEG. Journal of Neurophysiology, 91(6), 2649–2657.

Johns, M. W. (1991). A new method for measuring daytime sleepiness: the Epworth sleepiness scale. Sleep, 14(6), 540–545.

Ju, Y. E. S., Ooms, S. J., Sutphen, C., Macauley, S. L., Zangrilli, M. A., Jerome, G., Fagan, A. M., Mignot, E., Zempel, J. M., Claassen, J. A. H. R., & Holtzman, D. M. (2017). Slow wave sleep disruption increases cerebrospinal fluid amyloid-β levels. Brain, 140(8), 2104–2111. 10.1093/brain/awx148

Kirchner, W. K. (1958). Age differences in short-term retention of rapidly changing information. Journal of Experimental Psychology, 55(4), 352.

Klunk, W. E., Koeppe, R. A., Price, J. C., Benzinger, T. L., Devous, M. D., Jagust, W. J., Johnson, K. A., Mathis, C. A., Minhas, D., Pontecorvo, M. J., Rowe, C. C., Skovronsky, D. M., & Mintun, M. A. (2015). The Centiloid project: Standardizing quantitative amyloid plaque estimation by PET. Alzheimer’s and Dementia, 11(1), 1–15.e4. 10.1016/j.jalz.2014.07.003

Koch, S., Schmidtke, J., Krawczak, M., & Caliebe, A. (2023). Clinical utility of polygenic risk scores: a critical 2023 appraisal. Journal of Community Genetics, 14(5), 471–487.

Koshmanova, E., Muto, V., Chylinski, D., Mouraux, C., Reyt, M., Grinard, M., Talwar, P., Lambot, E., Berthomier, C., & Brandewinder, M. (2022). Genetic risk for insomnia is associated with objective sleep measures in young and healthy good sleepers. Neurobiology of Disease, 175, 105924.

Krasny, S., Yan, C., Hartley, S. L., Handen, B. L., Wisch, J. K., Boehrwinkle, A. H., Ances, B. M., Rafii, M. S., & consortium, A. (2024). Assessing amyloid PET positivity and cognitive function in Down syndrome to guide clinical trials targeting amyloid. Alzheimer’s & Dementia, 20(8), 5570–5577.

Leonenko, G., Shoai, M., Bellou, E., Sims, R., Williams, J., Hardy, J., Escott-Price, V., & Initiative, A. D. N. (2019). Genetic risk for alzheimer disease is distinct from genetic risk for amyloid deposition. Annals of Neurology, 86(3), 427–435.

Li, J., Vitiello, M. V, & Gooneratne, N. S. (2022). Sleep in normal aging. Sleep Medicine Clinics, 17(2), 161–171.

Lim, A. S. P., Kowgier, M., Yu, L., Buchman, A. S., & Bennett, D. A. (2013). Sleep Fragmentation and the Risk of Incident Alzheimer’s Disease and Cognitive Decline in Older Persons. Sleep, 36(7), 1027–1032. 10.5665/sleep.2802

Luckett, E. S., Abakkouy, Y., Reinartz, M., Adamczuk, K., Schaeverbeke, J., Verstockt, S., De Meyer, S., Van Laere, K., Dupont, P., & Cleynen, I. (2022). Association of Alzheimer’s disease polygenic risk scores with amyloid accumulation in cognitively intact older adults. Alzheimer’s Research & Therapy, 14(1), 138.

Ly, J. Q. M., Gaggioni, G., Chellappa, S. L., Papachilleos, S., Brzozowski, A., Borsu, C., Rosanova, M., Sarasso, S., Middleton, B., & Luxen, A. (2016). Circadian regulation of human cortical excitability. Nature Communications, 7(1), 11828.

Mahowald, M. W., & Schenck, C. H. (2005). Insights from studying human sleep disorders. Nature, 437(7063), 1279–1285.

Marks, S. M., Lockhart, S. N., Baker, S. L., & Jagust, W. J. (2017). Tau and β-amyloid are associated with medial temporal lobe structure, function, and memory encoding in normal aging. Journal of Neuroscience, 37(12), 3192–3201.

Mascetti, L., Foret, A., Schrouff, J., Muto, V., Dideberg, V., Balteau, E., Degueldre, C., Phillips, C., Luxen, A., & Collette, F. (2013). Concurrent synaptic and systems memory consolidation during sleep. Journal of Neuroscience, 33(24), 10182–10190.

Murphy, A. E., Schilder, B. M., & Skene, N. G. (2021). MungeSumstats: a Bioconductor package for the standardization and quality control of many GWAS summary statistics. Bioinformatics, 37(23), 4593–4596.

Muto, V., Jaspar, M., Meyer, C., Kussé, C., Chellappa, S. L., Degueldre, C., Balteau, E., Shaffii-Le Bourdiec, A., Luxen, A., & Middleton, B. (2016). Local modulation of human brain responses by circadian rhythmicity and sleep debt. Science, 353(6300), 687–690.

Muto, V., Koshmanova, E., Ghaemmaghami, P., Jaspar, M., Meyer, C., Elansary, M., Van Egroo, M., Chylinski, D., Berthomier, C., Brandewinder, M., Mouraux, C., Schmidt, C., Hammad, G., Coppieters, W., Ahariz, N., Degueldre, C., Luxen, A., Salmon, E., Phillips, C., … Vandewalle, G. (2021). Alzheimer’s disease genetic risk and sleep phenotypes in healthy young men: Association with more slow waves and daytime sleepiness. Sleep, 44(1), 1–12. 10.1093/sleep/zsaa137

Osorio-Forero, A., Cherrad, N., Banterle, L., Fernandez, L. M. J., & Lüthi, A. (2022). When the locus coeruleus speaks up in sleep: recent insights, emerging perspectives. International Journal of Molecular Sciences, 23(9), 5028.

Osorio-Forero, A., Foustoukos, G., Cardis, R., Cherrad, N., Devenoges, C., Fernandez, L. M. J., & Lüthi, A. (2025). Infraslow noradrenergic locus coeruleus activity fluctuations are gatekeepers of the NREM–REM sleep cycle. Nature Neuroscience, 28(1), 84–96.

Peter-Derex, L., Berthomier, C., Taillard, J., Berthomier, P., Bouet, R., Mattout, J., Brandewinder, M., & Bastuji, H. (2021). Automatic analysis of single-channel sleep EEG in a large spectrum of sleep disorders. Journal of Clinical Sleep Medicine, 17(3), 393–402.

Purcell, S., Neale, B., Todd-Brown, K., Thomas, L., Ferreira, M. A. R., Bender, D., Maller, J., Sklar, P., De Bakker, P. I. W., & Daly, M. J. (2007). PLINK: a tool set for whole-genome association and population-based linkage analyses. The American Journal of Human Genetics, 81(3), 559–575.

Reitz, C., Pericak-Vance, M. A., Foroud, T., & Mayeux, R. (2023). A global view of the genetic basis of Alzheimer disease. Nature Reviews Neurology, 19(5), 261–277.

Rigby, R. A., & Stasinopoulos, D. M. (2005). Generalized additive models for location, scale and shape. Journal of the Royal Statistical Society Series C: Applied Statistics, 54(3), 507–554.

Rigby, R. A., & Stasinopoulos, D. M. (2014). Automatic smoothing parameter selection in GAMLSS with an application to centile estimation. Statistical Methods in Medical Research, 23(4), 318–332.

Rigby, R. A., Stasinopoulos, M. D., Heller, G. Z., & De Bastiani, F. (2019). Distributions for modeling location, scale, and shape: Using GAMLSS in R. Chapman and Hall/CRC.

Roehrs, T., Merlotti, L., Petrucelli, N., Stepanski, E., & Roth, T. (1994). Experimental sleep fragmentation. Sleep, 17(5), 438–443.

Schmidt, C., Peigneux, P., & Cajochen, C. (2012). Age-related changes in sleep and circadian rhythms: Impact on cognitive performance and underlying neuroanatomical networks. Frontiers in Neurology, JUL(July), 1–11. 10.3389/fneur.2012.00118

Sharon, O., Zhelezniakov, V., Gat, Y., Falach, R., Narbayev, D., Shiner, T., Walker, M. P., Tauman, R., Bregman, N., & Nir, Y. (2025). Slow wave synchrony during NREM sleep tracks cognitive impairment in prodromal Alzheimer’s disease. Alzheimer’s & Dementia, 21(5), e70247.

Shi, L., Chen, S.-J., Ma, M.-Y., Bao, Y.-P., Han, Y., Wang, Y.-M., Shi, J., Vitiello, M. V, & Lu, L. (2018). Sleep disturbances increase the risk of dementia: a systematic review and meta-analysis. Sleep Medicine Reviews, 40, 4–16.

Stark, S. M., Yassa, M. A., Lacy, J. W., & Stark, C. E. L. (2013). A task to assess behavioral pattern separation (BPS) in humans: Data from healthy aging and mild cognitive impairment. Neuropsychologia, 51(12), 2442–2449.

Stasinopoulos, D. M., & Rigby, R. A. (2008). Generalized additive models for location scale and shape (GAMLSS) in R. Journal of Statistical Software, 23, 1–46.

Stasinopoulos, M. D., Rigby, R. A., Heller, G. Z., Voudouris, V., & De Bastiani, F. (2017). Flexible regression and smoothing: using GAMLSS in R. CRC Press, Taylor & Francis Group.

Steriade, M. (2003). The corticothalamic system in sleep. Front Biosci, 8(4), d878–d899.

Tabelow, K., Balteau, E., Ashburner, J., Callaghan, M. F., Draganski, B., Helms, G., Kherif, F., Leutritz, T., Lutti, A., & Phillips, C. (2019). hMRI–A toolbox for quantitative MRI in neuroscience and clinical research. Neuroimage, 194, 191–210.

Talwar, P., Sinha, J., Grover, S., Rawat, C., Kushwaha, S., Agarwal, R., Taneja, V., & Kukreti, R. (2016). Dissecting complex and multifactorial nature of Alzheimer’s disease pathogenesis: a clinical, genomic, and systems biology perspective. Molecular Neurobiology, 53(7), 4833–4864.

Tsai, C.-Y., Wu, S.-M., Kuan, Y.-C., Lin, Y.-T., Hsu, C.-R., Hsu, W.-H., Liu, Y.-S., Majumdar, A., Stettler, M., & Yang, C.-M. (2022). Associations between risk of Alzheimer’s disease and obstructive sleep apnea, intermittent hypoxia, and arousal responses: A pilot study. Frontiers in Neurology, 13, 1038735.

Tsapanou, A., Gao, Y., Stern, Y., & Barral, S. (2020). Polygenic score for sleep duration. Association with cognition. Sleep Medicine, 74, 262–266.

Tulsky, D., Zhu, J., & Ledbetter, M. F. (1997). WAIS-III/WMS-III technical manual. *Psychological Corporation*: *San Antonio*, *TX*.

Tzourio-Mazoyer, N., Landeau, B., Papathanassiou, D., Crivello, F., Etard, O., Delcroix, N., Mazoyer, B., & Joliot, M. (2002). Automated anatomical labeling of activations in SPM using a macroscopic anatomical parcellation of the MNI MRI single-subject brain. Neuroimage, 15(1), 273–289.

Van Egroo, M., Narbutas, J., Chylinski, D., Villar González, P., Maquet, P., Salmon, E., Bastin, C., Collette, F., & Vandewalle, G. (2019). Sleep–wake regulation and the hallmarks of the pathogenesis of Alzheimer’s disease. Sleep, 42(4). 10.1093/sleep/zsz017

Van Egroo, M., van Someren, E. J. W., Grinberg, L. T., Bennett, D. A., & Jacobs, H. I. L. (2024). Associations of 24-Hour Rest-Activity Rhythm Fragmentation, Cognitive Decline, and Postmortem Locus Coeruleus Hypopigmentation in Alzheimer’s Disease. Annals of Neurology.

Vandewalle, G., Archer, S. N., Wuillaume, C., Balteau, E., Degueldre, C., Luxen, A., Maquet, P., & Dijk, D.-J. (2009). Functional magnetic resonance imaging-assessed brain responses during an executive task depend on interaction of sleep homeostasis, circadian phase, and PER3 genotype. Journal of Neuroscience, 29(25), 7948–7956.

Wechsler, D. (2001). Echelle clinique de mémoire de Wechsler (MEM-III).

Weiskopf, N., & Helms, G. (2008). Multi-parameter mapping of the human brain at 1mm resolution in less than 20 minutes. Proceedings of ISMRM, 16, 2241.

Wightman, D. P., Jansen, I. E., Savage, J. E., Shadrin, A. A., Bahrami, S., Holland, D., Rongve, A., Børte, S., Winsvold, B. S., & Drange, O. K. (2021). A genome-wide association study with 1,126,563 individuals identifies new risk loci for Alzheimer’s disease. Nature Genetics, 53(9), 1276–1282.

Xicota, L., Gyorgy, B., Grenier-Boley, B., Lecoeur, A., Fontaine, G., Danjou, F., Gonzalez, J. S., Colliot, O., Amouyel, P., & Martin, G. (2022). Association of APOE-Independent Alzheimer disease polygenic risk score with brain amyloid deposition in asymptomatic older adults. Neurology, 99(5), e462–e475.

Zhang, Ren, R., Yang, L., Zhang, H., Shi, Y., Okhravi, H. R., Vitiello, M. V, Sanford, L. D., & Tang, X. (2022). Sleep in Alzheimer’s disease: a systematic review and meta-analysis of polysomnographic findings. Translational Psychiatry, 12(1), 136.

Zou, G., Xu, J., Zhou, S., Liu, J., Su, Z. H., Zou, Q., & Gao, J.-H. (2020). Functional MRI of arousals in nonrapid eye movement sleep. Sleep, 43(2), zsz218.

