## supplementary file for "Sleep arousals are associated with the polygenic risk for developing Alzheimer’s disease and with cognitive decline in healthy late middle-aged individuals"

**Supplementary Table S1. Association between arousals number and PRS for AD after adding the variables that significantly differed between the two age groups (i.e., BMI, depression, sleep quality and chronotype) as covariates.**

| Independent variable | t | Estimate | SE | p-value |
|---|---|---|---|---|
| **PRS** | -1.05 | -0.005 | 0.005 | 0.29 |
| **Transition status** | 41.14 | 1.30 | 0.03 | **<.0001** |
| **EMG status** | -0.83 | -0.03 | 0.04 | 0.41 |
| **Age group** | -8.27 | -0.29 | 0.03 | **<.0001** |
| **Sex** | 2.68 | 0.05 | 0.02 | **<.0001** |
| **TST** | 38.64 | 0.003 | 0.00007 | **<.0001** |
| **BMI** | -7.90 | -0.01 | 0.001 | **<.0001** |
| **Depression** | -5.32 | -0.005 | 0.0009 | **<.0001** |
| **Chronotype** | 6.13 | 0.002 | 0.0004 | **<.0001** |
| **Sleep quality** | 2.36 | 0.004 | 0.002 | **<.0001** |
| **PRS*Transition** | 1.66 | 0.01 | 0.006 | 0.10 |
| **PRS*EMG** | 1.69 | 0.01 | 0.007 | 0.09 |
| **Transition*EMG** | 13.16 | 0.58 | 0.04 | **<.0001** |
| **PRS* age group** | 0.79 | 0.005 | 0.006 | 0.43 |
| **Transition * age group** | 22.58 | 0.80 | 0.04 | **<.0001** |
| **EMG*age group** | -6.91 | -0.32 | 0.05 | **<.0001** |
| **PRS*Transition*EMG** | -2.40 | -0.02 | 0.008 | **<.0001** |
| **PRS*Transition*age group** | -1.54 | -0.01 | 0.006 | 0.12 |
| **PRS*EMG*age group** | -1.36 | -0.01 | 0.008 | 0.17 |
| **Transition*EMG*age group** | -1.41 | -0.07 | 0.05 | 0.16 |
| **PRS*Transition*EMG* age group** | 2.57 | 0.02 | 0.009 | **<.0001** |

Analyses were performed using a generalized additive model for location, scale, and shape (GAMLSS). Results with p values less than 0.05 were considered statistically significant (df=1447.663).

BMI: Body mass index; PRS: polygenic risk score; TST: total sleep time; EMG: Electromyography.

This analysis shows that despite significant associations with the additional variable, the main outcome of the GAMLSS remains unchanged.

**Supplementary Table S2. Association between sex and T+M+ and T+M− arousals in young and late middle-aged individuals.**

| Age group | Arousal type (dependent variable) | PRS | PRS*Sex | Sex | Age | Total sleep time |
|---|---|---|---|---|---|---|
| **Young individuals** | **T+M+ arousals number (N=452)** | t(444)=0.68 Estimate=0.03 SE=0.04 p=0.50 | t(444)=-0.93 Estimate=-0.12 SE=0.13 p=0.36 | t(444)=0.66 Estimate=0.51 SE=0.78 p=0.51 | t(444)=3.55 Estimate=0.28 SE=0.08 p=0.0004 | t(444)=5.79 Estimate=0.03 SE=0.005 p=0.00000001 |
| | **T+M- arousals number (N=449)** | t(441)=0.10 Estimate=0.004 SE=0.04 p=0.92 | t(441)=-0.16 Estimate=-0.02 SE=0.12 p=0.88 | t(441)=3.92 Estimate=2.19 SE=0.56 p=0.0001 | t(441)=0.80 Estimate=0.05 SE=0.06 p=0.42 | t(441)=3.45 Estimate=0.02 SE=0.005 p=0.0006 |
| **Late middle-aged individuals** | **T+M+ arousals number (N=87)** | t(79)=-2.07 Estimate=-0.03 SE=0.01 **p=0.04** | t(79)=1.83 Estimate=0.03 SE=0.02 p=0.07 | t(79)=-2.34 Estimate=-0.19 SE=0.08 p=0.02 | t(79)=2.15 Estimate=0.02 SE=0.007 p=0.03 | t(79)=2.10 Estimate=0.002 SE=0.0009 p=0.04 |
| | **T+M- arousals number (N=86)** | t(78)=0.91 Estimate=0.31 SE=0.34 p=0.37 | t(78)=-0.005 Estimate=-0.002 SE=0.37 p=0.99 | t(78)=1.90 Estimate=3.31 SE=1.74 p=0.06 | t(78)=1.21 Estimate=0.16 SE=0.13 p=0.23 | t(78)=0.76 Estimate=0.02 SE=0.02 p=0.45 |

Analyses were performed using a generalized additive model for location, scale, and shape (GAMLSS). Results with p values less than 0.05 were considered statistically significant. Estimates in GAMLSS are reported in the log scale.

Prior to the analysis, we removed the outliers among all variables by excluding the samples lying beyond four times the standard deviation (the final number of individuals included in each analysis is reported below each dependent variable).

The fact that we find no significant interaction between sex and PRS for AD suggest that the association between arousal types and PRS in late middle-aged individuals is not different between men and women. See also figure S2 and S3 for displays.

**Supplementary Table S3. Association between PRS for AD (as dependent variable) and arousals number in REM and NREM separately.**

| Independent variable | t | Estimate | SE | p-value |
|---|---|---|---|---|
| **T+M+ REM** | -0.75 | -0.31 | 0.41 | 0.45 |
| **T+M+ NREM** | -1.80 | -0.25 | 0.14 | 0.07 |
| **T-M+ REM** | 0.23 | 0.01 | 0.05 | 0.81 |
| **T-M+ NREM** | 1.26 | 0.05 | 0.04 | 0.21 |
| **T+M-NREM** | 2.51 | 0.28 | 0.11 | **0.01** |
| **T-M-NREM** | -1.29 | -0.03 | 0.02 | 0.20 |
| **T+M+ REM *group** | 0.62 | 0.28 | 0.45 | 0.53 |
| **T+M+ NREM *group** | 1.60 | 0.25 | 0.15 | 0.11 |
| **T-M+ REM *group** | -0.60 | -0.04 | 0.06 | 0.55 |
| **T-M+ NREM *group** | -0.98 | -0.04 | 0.04 | 0.33 |
| **T+M-NREM *group** | -2.28 | -0.29 | 0.13 | **0.02** |
| **T-M-NREM *group** | 1.46 | 0.03 | 0.02 | 0.15 |
| **group** | -0.24 | -0.61 | 2.50 | 0.81 |
| **sex** | -0.08 | -0.06 | 0.76 | 0.94 |
| **TST** | -0.006 | -0.00004 | 0.007 | 1.00 |

Analyses were performed using a generalized additive model for location, scale, and shape (GAMLSS). Results with p values less than 0.05 were considered statistically significant (df=515). Estimates in GAMLSS are reported in the log scale.

Prior to the analysis, we removed the outliers among all variables by excluding the samples lying beyond four times the standard deviation.

TST: total sleep time.

The fact that we find no association between PRS for AD and T+M+ arousals when separated between REM and NREM sleep suggests that it is the total number of T+M+ in REM and NREM that was associated with the PRS for (NB: according to ASSM criteria, arousals must be associated with muscle tone increase during REM, such that all M- arousal were only detected during NREM).

**Supplementary Table S4. Non-significant associations between cognitive domains at baseline and T+M+ and T+M− arousals in young individuals.**

| Cognitive domain (dependent variable) | T+M+ arousals number | T+M- arousals number | Age | Education | Total sleep time |
|---|---|---|---|---|---|
| **Attention (N=286)** | t(277)=-1.31 Estimate=-0.02 SE=0.02 p=0.19 | t(277)=-1.06 Estimate=-0.02 SE=0.02 p=0.29 | t(277)=-1.89 Estimate=-0.08 SE=0.04 p=0.06 | t(277)=2.64 Estimate=0.20 SE=0.08 p=0.009 | t(277)=-0.11 Estimate=-0.0003 SE=0.002 p=0.91 |
| **Memory (N=304)** | t(295)=-0.68 Estimate=-0.01 SE=0.02 p=0.50 | t(295)=-0.69 Estimate=-0.02 SE=0.02 p=0.49 | t(295)=0.20 Estimate=0.009 SE=0.05 p=0.84 | t(295)=0.34 Estimate=0.03 SE=0.08 p=0.74 | t(295)=-0.69 Estimate=-0.002 SE=0.003 p=0.49 |
| **Executive function (N=278)** | t(269)=-0.70 Estimate=-0.02 SE=0.02 p=0.48 | t(269)=-0.26 Estimate=-0.008 SE=0.03 p=0.79 | t(269)=-1.06 Estimate=-0.07 SE=0.07 p=0.29 | t(269)=3.06 Estimate=0.35 SE=0.11 p=0.002 | t(269)=-0.25 Estimate=-0.0008 SE=0.003 p=0.80 |

Analyses were performed using a generalized additive model for location, scale, and shape (GAMLSS). Results with p values less than 0.05 were considered statistically significant. Estimates in GAMLSS are reported in the log scale.

Prior to the analysis, we removed the outliers among all variables by excluding the samples lying beyond four times the standard deviation (the final number of individuals included in each analysis is reported below each dependent variable).

Of note, all individuals did not finish all cognitive tests.

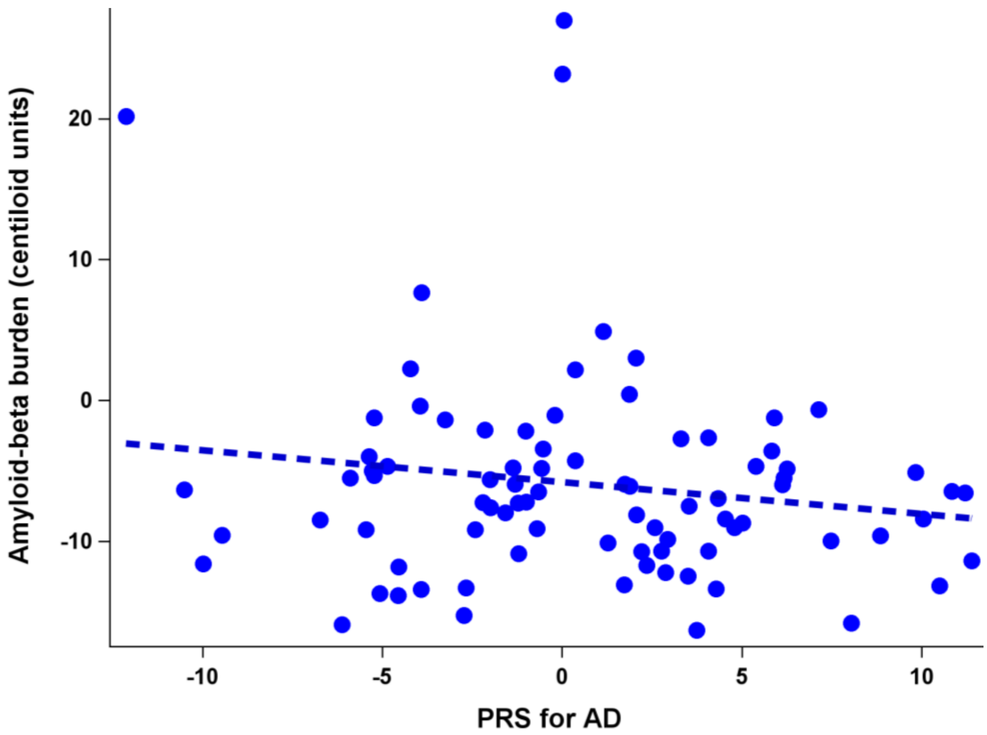


**Supplementary Figure S1.** Association between the polygenic risk score (PRS) for Alzheimer's disease and amyloid-beta (Aβ) burden including Aβ-positive individuals. Positivity was set at Aβ centiloid > 20 (Krasny et al., 2024) . Sperman's r=-.08; p=.43. The figure is complementary to Figure 2A.

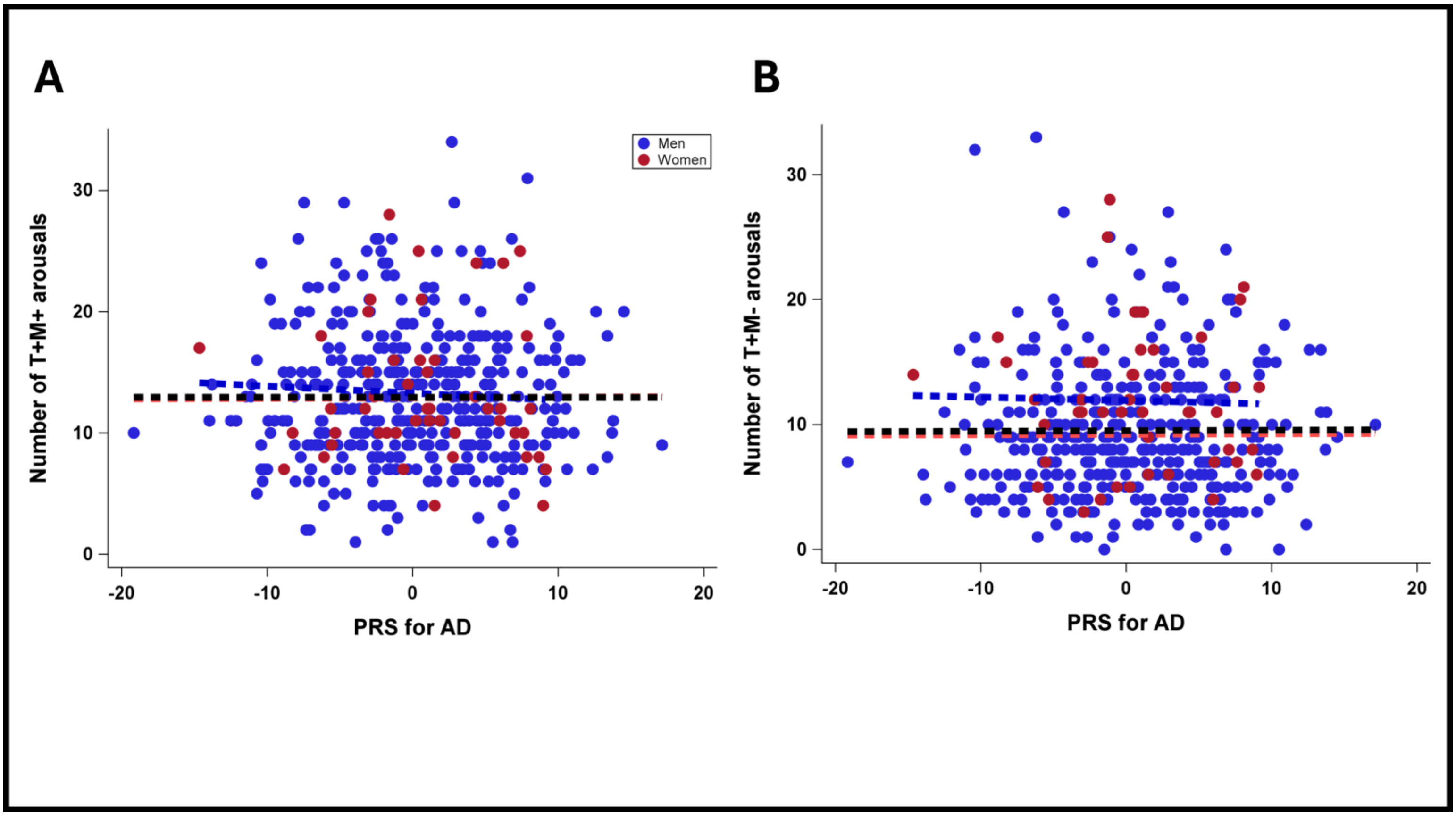


**Supplementary Figure S2. Non-significant associations between significant arousal types and the interaction of sex and PRS for AD in young individuals. (A)** Association between the number of T+M+ arousals and PRS for AD in young individuals (p=0.36). **(B)** Association between the number of T+M- arousals and PRS for AD (p=0.88).

Simple regression lines are used for a visual display and do not substitute the GAMLSS outputs (see Suppl. Table S2). The black line represents the regression irrespective of age groups (young + old). Dashed regression lines represent non-significant outputs of the GAMLSS.

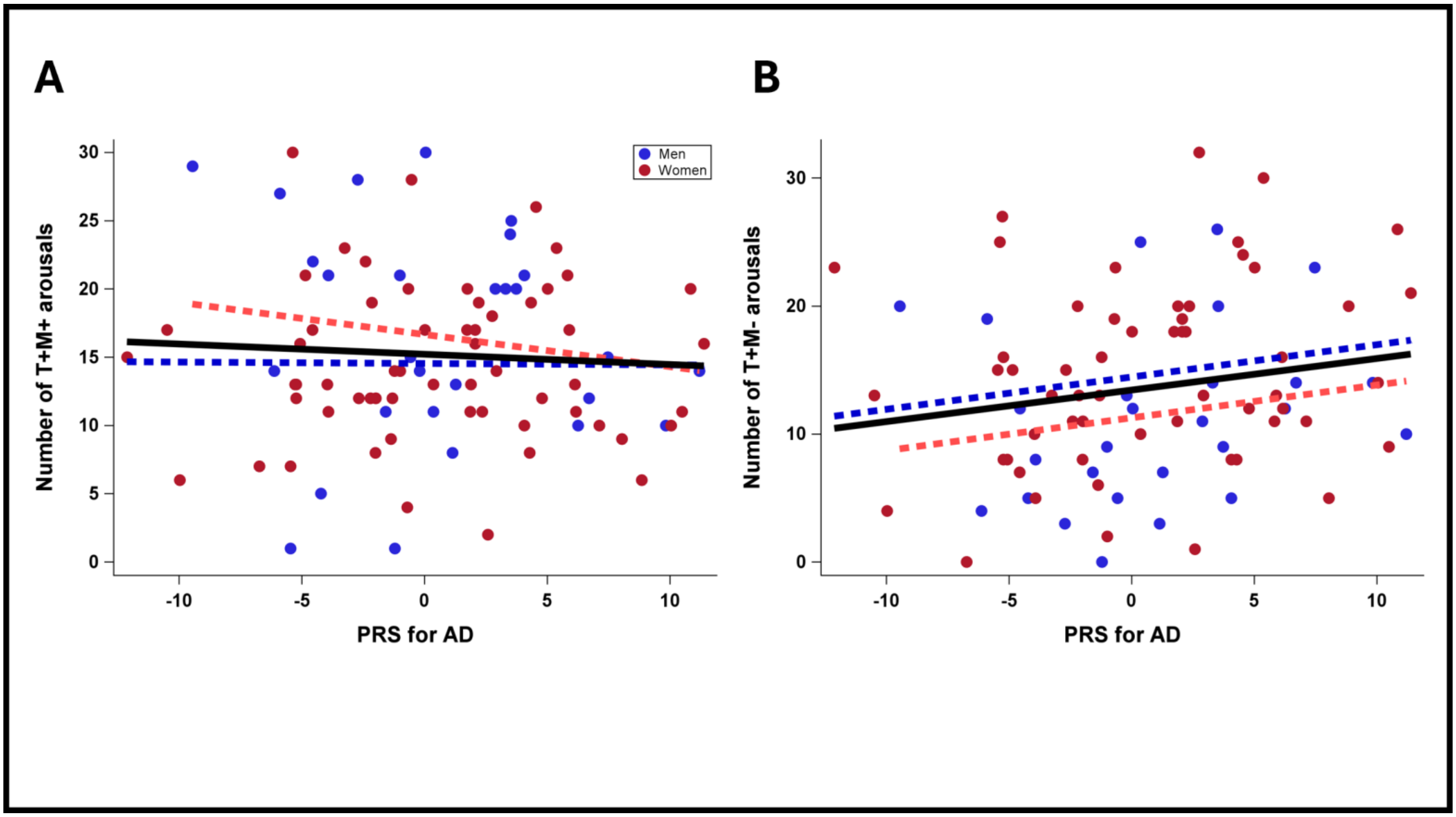


**Supplementary Figure S3. Non-significant associations between significant arousal types and the interaction of sex and PRS for AD in the late middle aged individuals. (A)** Association between the number of T+M+ arousals and PRS for AD (p=0.07). **(B)** Association between the number of T+M- arousals and PRS for AD (p=0.99).

Simple regression lines are used for a visual display and do not substitute the GAMLSS outputs (see Suppl. Table S2). The black line represents the regression irrespective of age groups (young + old). Solid and dashed regression lines represent significant and non-significant outputs of the GAMLSS, respectively.

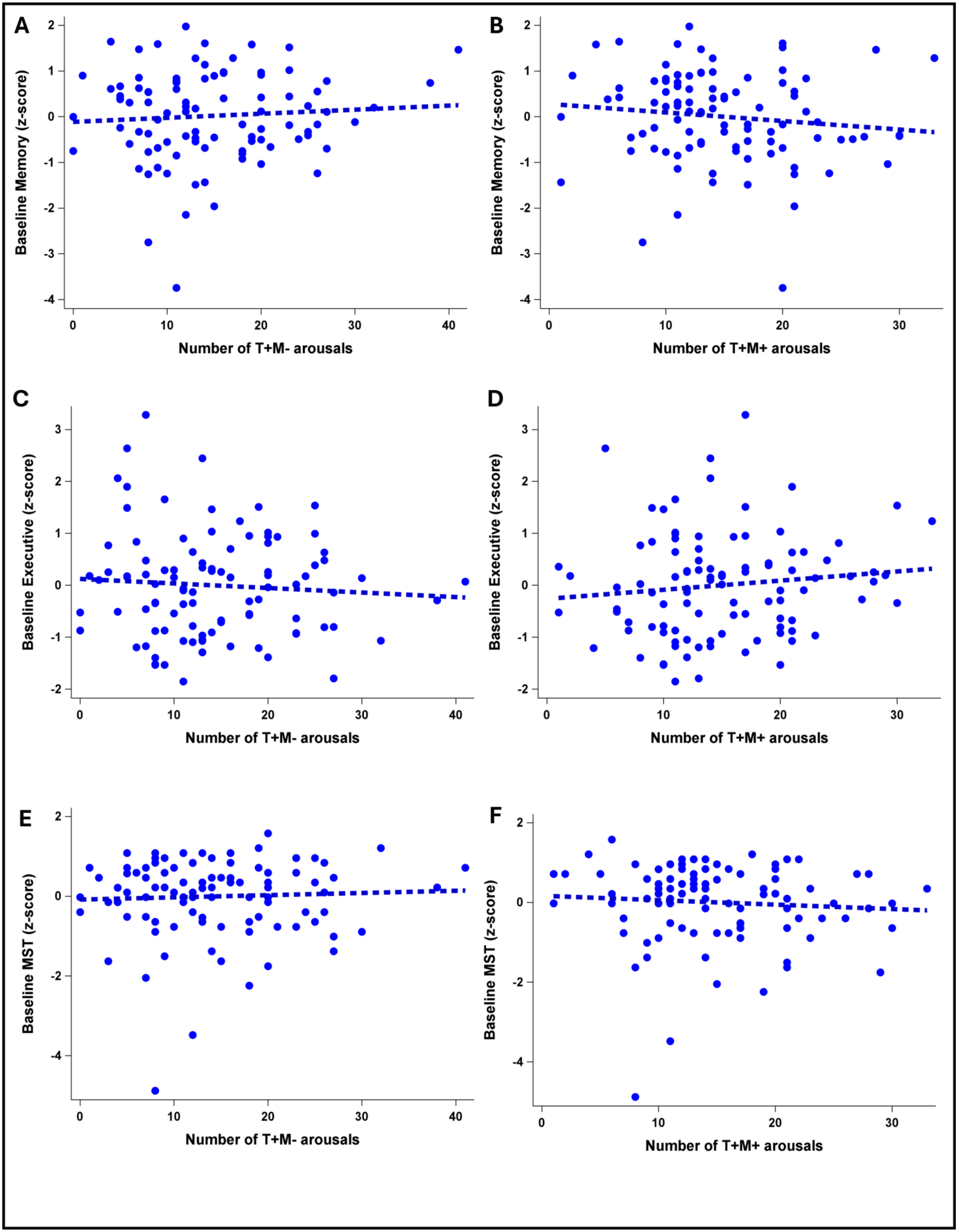

A
B
C
D
E
F
Baseline Memory (z-score)
Baseline Executive (z-score)
Baseline MST (z-score)
Number of T+M- arousals
Number of T+M+ arousals

**Supplementary Figure S4. Non-significant associations between baseline cognitive scores and T+M+ and T+M- arousals in late middle-aged individuals. (A)** Association between the number of T+M- arousals and baseline memory scores. **(B)** Association between the number of T+M+ arousals and baseline memory scores. **(C)** Association between the number of T+M- arousals and baseline executive scores. **(D)** Association between the number of T+M+ arousals and baseline executive scores. **(E)** Association between the number of T+M- arousals and baseline MST scores. **(F)** Association between the number of T+M+ arousals and baseline MST scores.

Simple regression lines are used for a visual display and do not substitute the GAMLSS outputs (see Table 4). The dashed regression lines represent non-significant outputs of the GAMLSS. None of the associations were significant ($p>0.06$).

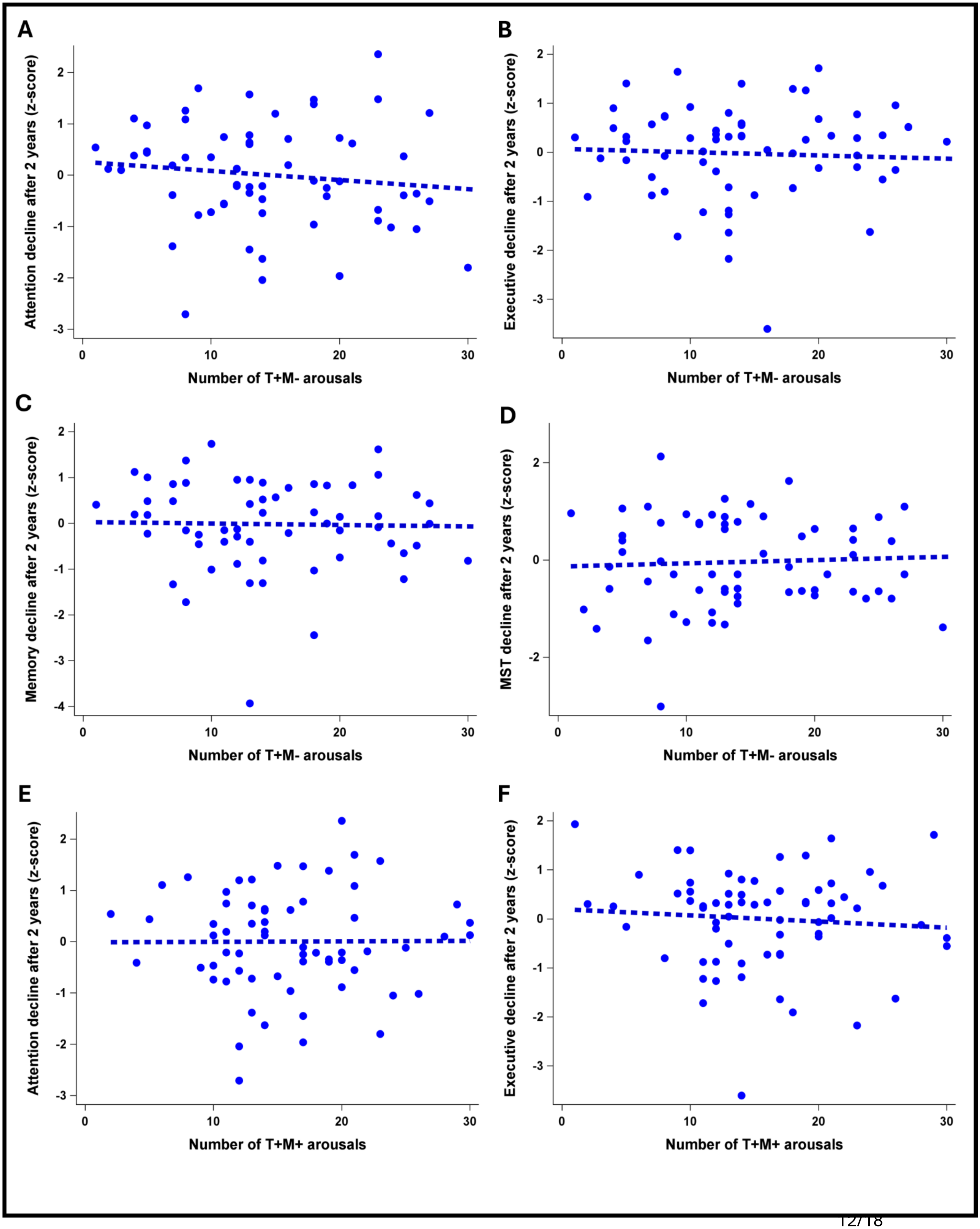
A
Attention decline after 2 years (z-score)
Number of T+M- arousals
B
Executive decline after 2 years (z-score)
Number of T+M- arousals
C
Memory decline after 2 years (z-score)
Number of T+M- arousals
D
MST decline after 2 years (z-score)
Number of T+M- arousals
E
Attention decline after 2 years (z-score)
Number of T+M+ arousals
F
Executive decline after 2 years (z-score)
Number of T+M+ arousals

**Supplementary Figure S5. Non-significant associations between cognitive decline after two years and T+M+ and T+M- arousals in late middle-aged individuals. (A)** Association between the number of T+M- arousals and attention decline after 2 years. **(B)** Association between the number of T+M- arousals and executive decline after 2 years. **(C)** Association between the number of T+M- arousals and memory decline after 2 years. **(D)** Association between the number of T+M- arousals and MST decline after 2 years. **(E)** Association between the number of T+M+ arousals and attention decline after 2 years. **(F)** Association between the number of T+M+ arousals and executive decline after 2 years.

Simple regression lines are used for a visual display and do not substitute the GAMLSS outputs (see Table 4). The dashed regression lines represent non-significant outputs of the GAMLSS. None of the associations were significant ($p>0.20$).

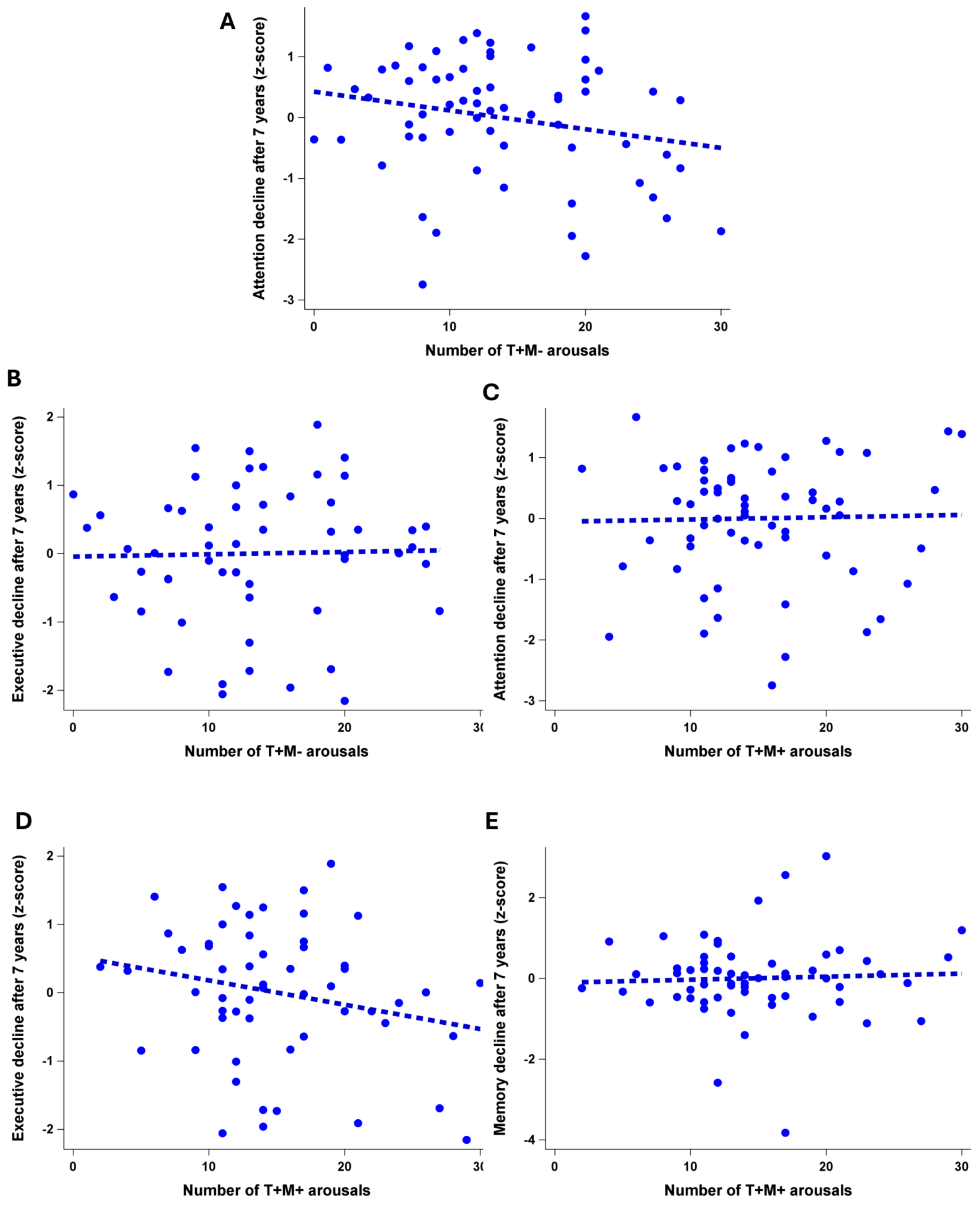
A
Attention decline after 7 years (z-score)
Number of T+M- arousals
B
Executive decline after 7 years (z-score)
Number of T+M- arousals
C
Attention decline after 7 years (z-score)
Number of T+M+ arousals
D
Executive decline after 7 years (z-score)
Number of T+M+ arousals
E
Memory decline after 7 years (z-score)
Number of T+M+ arousals

**Supplementary Figure S6. Non-significant associations between cognitive decline after seven years and T+M+ and T+M- arousals in late middle-aged individuals. (A)** Association between the number of T+M- arousals and attention decline after 7 years. **(B)** Association between the number of T+M- arousals and executive decline after 7 years. **(C)** Association between the number of T+M+ arousals and attention decline after 7 years. **(D)** Association between the number of T+M+ arousals and executive decline after 7 years. **(E)** Association between the number of T+M+ arousals and memory decline after 7 years.

None of the associations were significant ($p>0.10$). Simple regression lines are used for a visual display and do not substitute the GAMLSS outputs (see Table 4). The dashed regression lines represent non-significant outputs of the GAMLSS.

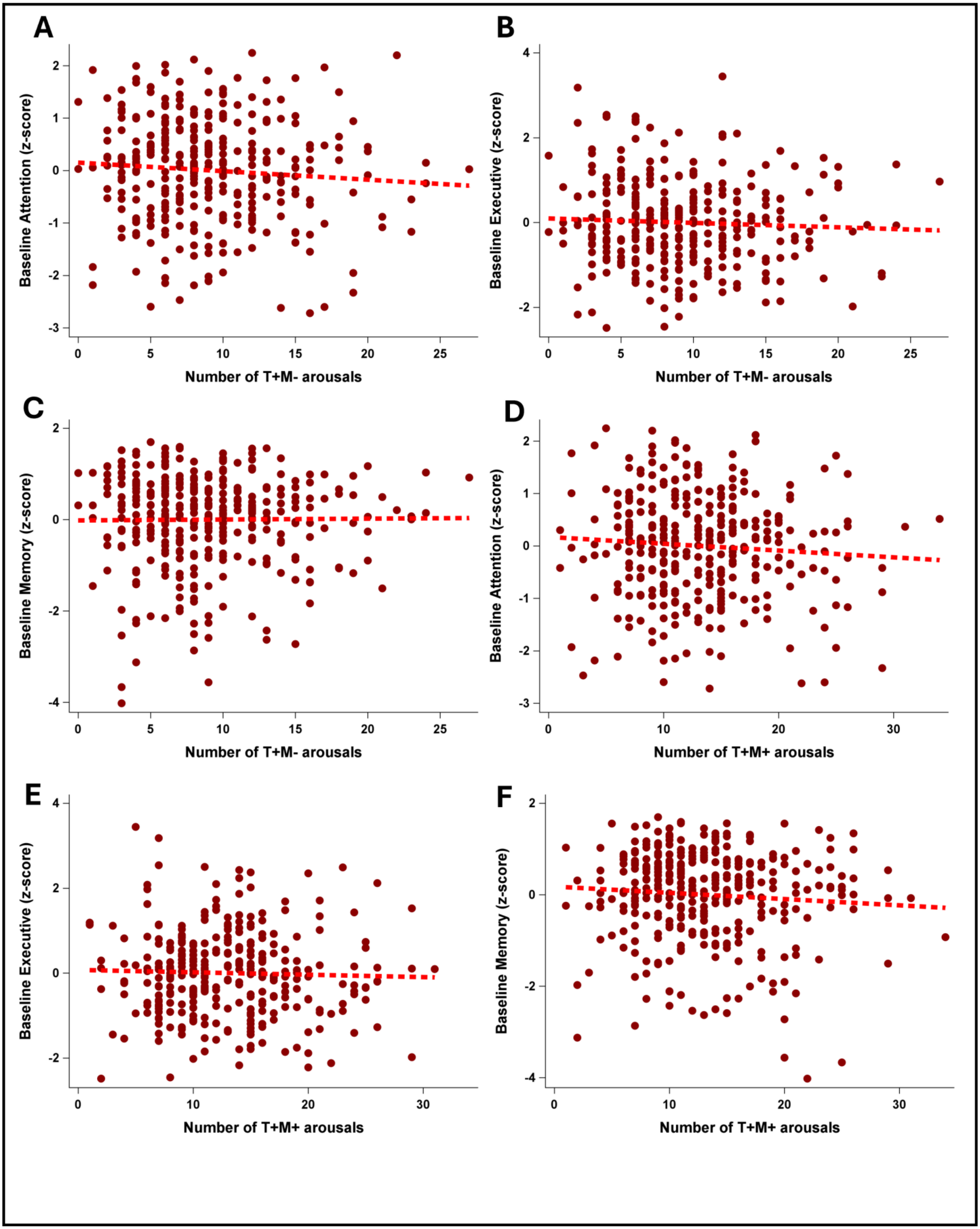

A
Baseline Attention (z-score)
Number of T+M- arousals
B
Baseline Executive (z-score)
Number of T+M- arousals
C
Baseline Memory (z-score)
Number of T+M- arousals
D
Baseline Attention (z-score)
Number of T+M+ arousals
E
Baseline Executive (z-score)
Number of T+M+ arousals
F
Baseline Memory (z-score)
Number of T+M+ arousals

**Supplementary Figure S7. Non-significant associations between baseline cognitive scores and T+M+ and T+M- arousals in young individuals. (A)** Association between the number of T+M- arousals and baseline attention scores. **(B)** Association between the number of T+M- arousals and baseline executive scores. **(C)** Association between the number of T+M- arousals and baseline memory scores. **(D)** Association between the number of T+M+ arousals and baseline attention scores. **(E)** Association between the number of T+M+ arousals and baseline executive scores. **(F)** Association between the number of T+M+ arousals and baseline memory scores.

None of the associations were significant ($p>0.19$). Refer to Suppl. Table S4 for detailed statistical outputs.

Domain-specific composites scores were computed for the memory, executive function, and attentional domains, and they consisted of the standardized (z- scores) sum of the standardized domain-specific scores, where higher values indicate better performance. The memory score consisted of the List of words I and II and family scenes II from Wechsler clinical scale of memory assessment MEM-III (Wechsler, 2001). The executive function score included verbal fluency tests (letter and animals score for 2 minutes) (Cardebat et al., 1990), inverse order digit span (Tulsky et al., 1997) and Stroop task (Stroop, 1935). The attentional score comprised the Paced auditory serial addition test (PASAT) (Gronwall, 1977) and D2 attention test (Brickenkamp, 1966).

**Supplement references**


Brickenkamp, R. (1966). *D2R—d2 Test of Attention*. Hogrefe.

Cardebat, D., Doyon, B., Puel, M., Goulet, P., & Joanette, Y. (1990). Formal and semantic lexical evocation in normal subjects. Performance and dynamics of production as a function of sex, age and educational level. *Acta Neurologica Belgica*, *90*(4), 207–217.

Gronwall, D. M. (1977). Paced auditory serial-addition task: a measure of recovery from concussion. *Perceptual and Motor Skills*, *44*(2), 367–373. https://doi.org/10.2466/PMS.1977.44.2.367

Krasny, S., Yan, C., Hartley, S. L., Handen, B. L., Wisch, J. K., Boehrwinkle, A. H., Ances, B. M., Rafii, M. S., & consortium, A. (2024). Assessing amyloid PET positivity and cognitive function in Down syndrome to guide clinical trials targeting amyloid. *Alzheimer's & Dementia*, *20*(8), 5570–5577.

Stroop, J. R. (1935). Studies of interference in serial verbal reactions. *Journal of Experimental Psychology*, *18*(6), 643.

Tulsky, D., Zhu, J., & Ledbetter, M. F. (1997). WAIS-III/WMS-III technical manual. *Psychological Corporation: San Antonio, TX*.

Wechsler, D. (2001). *MEM-III, Échelle clinique de mémoire de Wechsler : manuel (3e éd.) David Wechsler*.
